# Comparative analysis of primary auditory cortical responses in bats and mice to repetitive stimuli trains

**DOI:** 10.1101/2022.10.28.514155

**Authors:** Katrina E. Deane, Francisco García-Rosales, Ruslan Klymentiev, Julio C. Hechavarria, Max F. K. Happel

## Abstract

The brains of black 6 mice (Mus musculus) and Seba’s short-tailed bats (Carollia perspicillata) weigh roughly the same and share mammalian neocortical laminar architecture. Bats have highly developed sonar calls and social communication and are an excellent neuroethological animal model for auditory research. Mice are olfactory and somatosensory specialists, used frequently in auditory neuroscience for their advantage of standardization and wide genetic toolkit. This study presents an analytical approach to overcome the challenge of inter-species comparison with existing data. In both data sets, we recorded with linear multichannel electrodes down the depth of the primary auditory cortex (A1) while presenting repetitive stimuli trains at ~5 and ~40 Hz to awake bats and mice. We found that while there are similarities between cortical response profiles in both, there was a better signal to noise ratio in bats under these conditions, which allowed for a clearer following response to stimuli trains. Model fit analysis supported this, illustrating that bats had stronger response amplitude suppression to consecutive stimuli. Additionally, continuous wavelet transform revealed that bats had significantly stronger power and phase coherence during stimulus response and mice had stronger power in the background. Better signal to noise ratio and lower intertrial phase variability in bats could represent specialization for faster and more accurate temporal processing at lower metabolic costs. Our findings demonstrate a potentially different general auditory processing principle; investigating such differences may increase our understanding of how the ecological need of a species shapes the development and function of its nervous system.

## Introduction

The brains and bodies of black 6 mice (*Mus musculus)* and Seba’s short-tailed bats (*Carollia perspicillata)* weigh roughly the same. Bats make up the second largest extant mammalian order, *Chiroptera*, after rodents, *Rodentia*, and they are the only mammals that can achieve true flight—converging their evolution with birds. Instead of sharing brain architecture with other flyers, they share neocortical laminar structures and microcircuitry with the rest of mammals, such as mice and humans (Chang & Kawai, 2018; García-Rosales et al., 2019; Linden & Schreiner, 2003; Mountcastle, 1997). Bats have highly developed sonar calls and social communication (Beetz et al., 2017; Hechavarría, Macías, Vater, Mora, et al., 2013; Thies et al., 1998; Weineck et al., 2020), making them a choice target for neuroethological auditory studies. Mice have a smaller repertoire of social verbal cues (Fonseca et al., 2021) and rely most heavily on their whiskers and olfaction for navigation (Gire et al., 2016), but they are frequently used in auditory neuroscience due to standardization and a wide transgenic toolkit.

There are two primary motivations for an exploratory cross-species analysis of the primary auditory cortex. First, investigating the potentially different application of general auditory processing would help to gain a better understanding of how the ecological needs of a species shape the development and function of the nervous system. Second, studies of the A1 may list a variety of species in their literature review to exemplify peer-reviewed findings, such as about oscillatory activity (e.g. cortical gamma in **mouse**: Chen et al., 2017: Shahriari et al., 2016; in **rat**: MacDonald & Barth, 1995: Vianney-Rodrigues et al., 2011; in **cat**: Karmos et al., 2002; Lakatos et al., 2004; etc.) or cortical layer roles (e.g. in **bat**: García-Rosales et al., 2019; in **mouse**: Chang & Kawai, 2018; in **rabbit**: McMullen & Glaser, 1982; in **cat**: Winguth & Winer, 1986, in **primate** Hashikawa et al., 1995; etc.). However, the ecology and evolutionary biology of the model may lead to A1 discrepancies in a species-specific way that has not been previously quantifiable through comparing across publications nor, to the authors’ knowledge, reviewed on a broad scale. While there are some studies comparing the A1 and dorsal auditory cortical areas of several bat species (e.g. Hagemann et al., 2011; Hechavarría et al., 2013), few compare the A1 of bats to other mammals (see Kanwal & Rauschecker, 2007). Similarly, there are few studies quantitatively comparing the mouse A1 against other species (see Hoglen et al., 2018). Generally, the functional differences between small mammals’ auditory abilities are open for exploration.

In this paper, we investigated the A1 of awake, head-fixed, freely moving black 6 mice and awake, head-fixed Seba’s short-tailed bats. We evaluated two existing A1 multichannel datasets across both species to perform comparative analyses aimed at understanding fundamental auditory response profiles between them. We hypothesized that bats would have more temporally accurate auditory processing than mice, due to their experience with echolocation and social communication. However, it was unclear how this would be achieved, given that both species share mammalian cortical architecture. Bats listened to a repeated distress syllable at 5.28 and 36.76 Hz and mice listened to click trains at 5 and 40 Hz, creating a possibly different valence in stimulus interpretation. Nevertheless, we explored laminar profiles with current source density (CSD) analysis and found discrepancies we believe to be ecologically founded beyond stimulus difference. We performed a model fit analysis to better understand temporal response and background suppression over consecutively repeated stimuli trains across these two data sets. We further ran continuous wavelet transform (CWT) analysis to compare internal coherence dynamics and signal to noise ratio differences of normalized spectral power. We also computed phase amplitude coupling (PAC) to investigate possible contributions to information transfer and spectral coupling profiles.

Overall, we found that the laminar flow of cortical activity in response to stimuli was highly conserved across short-tailed bats and mice. However, bats demonstrated a better signal to noise ratio under these conditions, meaning a stronger stimulus response to relative background noise. This was demonstrated in AVREC and layer traces and a subsequent model fit analysis, showing more robust stimulus-related activity, more accurate temporal resolution in response to stimuli trains, and higher suppression of consecutive stimuli. CWT analysis also revealed a stronger broadband oscillatory frequency power distribution during stimulus response, relative to the background, and stronger phase coherence during stimulus response in bats. PAC profiles were fundamentally different between species, with bats having stimulus specific coupling profile difference compared to mice. While the two studies under investigation differed in some methodological and experimental conditions, the statistical analyses provide potentially meaningful insight into the divergent recruitment of shared mammalian auditory physiology, linking evolutionary and behavioral need to specific auditory ability.

## Methods

Each dataset from the bat and mouse groups was collected for separate publication (see Table 1). Table 1 describes the crucial differences between data collection and parameters. All analyses in comparison of these two datasets are novel.

**Table 1.**
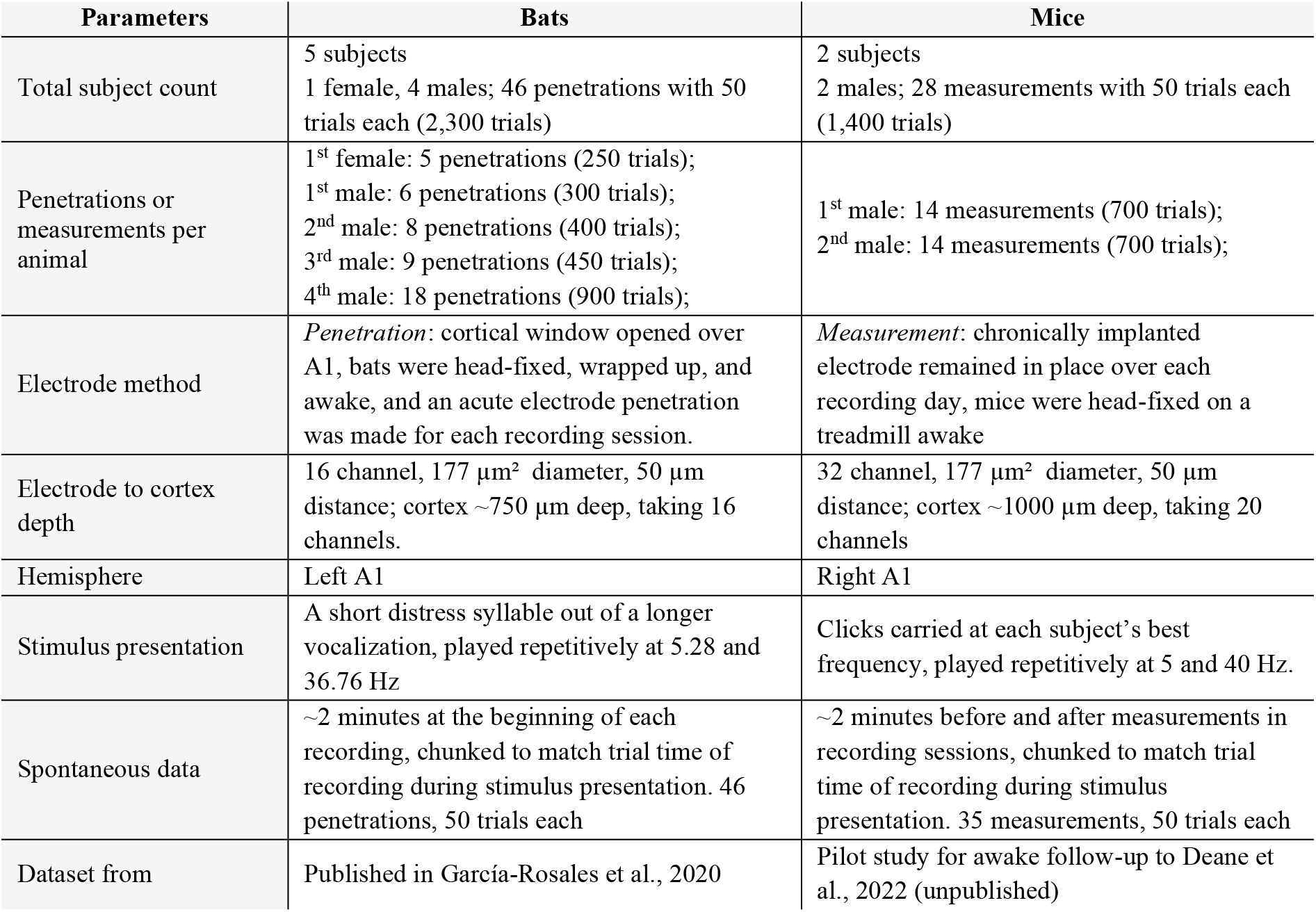
Parameter Table describing number of subjects, how many penetrations/measurements and trials were taken per subject, electrode method (briefly), electrode depth to cortex, hemisphere, description of stimulus and spontaneous recordings, and dataset source for each group

| Parameters | Bats | Mice |
| --- | --- | --- |
| Total subject count | 5 subjects<br>1 female, 4 males; 46 penetrations with 50 trials each (2,300 trials) | 2 subjects<br>2 males; 28 measurements with 50 trials each (1,400 trials) |
| Penetrations or measurements per animal | 1<sup>st</sup> female: 5 penetrations (250 trials);<br>1<sup>st</sup> male: 6 penetrations (300 trials);<br>2<sup>nd</sup> male: 8 penetrations (400 trials);<br>3<sup>rd</sup> male: 9 penetrations (450 trials);<br>4<sup>th</sup> male: 18 penetrations (900 trials); | 1<sup>st</sup> male: 14 measurements (700 trials);<br>2<sup>nd</sup> male: 14 measurements (700 trials); |
| Electrode method | *Penetration*: cortical window opened over A1, bats were head-fixed, wrapped up, and awake, and an acute electrode penetration was made for each recording session. | *Measurement*: chronically implanted electrode remained in place over each recording day, mice were head-fixed on a treadmill awake |
| Electrode to cortex depth | 16 channel, 177 $\mu\text{m}^2$ diameter, 50 $\mu\text{m}$ distance; cortex $\sim 750 \mu\text{m}$ deep, taking 16 channels. | 32 channel, 177 $\mu\text{m}^2$ diameter, 50 $\mu\text{m}$ distance; cortex $\sim 1000 \mu\text{m}$ deep, taking 20 channels |
| Hemisphere | Left A1 | Right A1 |
| Stimulus presentation | A short distress syllable out of a longer vocalization, played repetitively at 5.28 and 36.76 Hz | Clicks carried at each subject's best frequency, played repetitively at 5 and 40 Hz. |
| Spontaneous data | $\sim 2$ minutes at the beginning of each recording, chunked to match trial time of recording during stimulus presentation. 46 penetrations, 50 trials each | $\sim 2$ minutes before and after measurements in recording sessions, chunked to match trial time of recording during stimulus presentation. 35 measurements, 50 trials each |
| Dataset from | Published in García-Rosales et al., 2020 | Pilot study for awake follow-up to Deane et al., 2022 (unpublished) |

### Ethical Approval

All experiments were conducted in accordance with ethical animal research standards defined by the German Law and approved by an ethics committee of the State of Saxony-Anhalt under the given license 42502-2-1394LIN for mice and by the Darmstadt Regional Counsel under the given license #FU-1126 for bats. They also conform to the principles and regulations as described in by Grundy (2015). All experiments were carried out with adult male C57/B6 mice (*Mus musculus*, n = 2, 11-13 weeks of age, 24-28 g body weight) and adult Seba’s short-tailed bats (*Carollia perspicillata*, n = 5, 18-20 g body weight). Note that female mice were not used and 1 female bat (subject 1 in Supp Figure 1 and Supp Figure 2) was included in the bat dataset; possible variances due to sex was not in the scope of our study.

**Figure 1.**
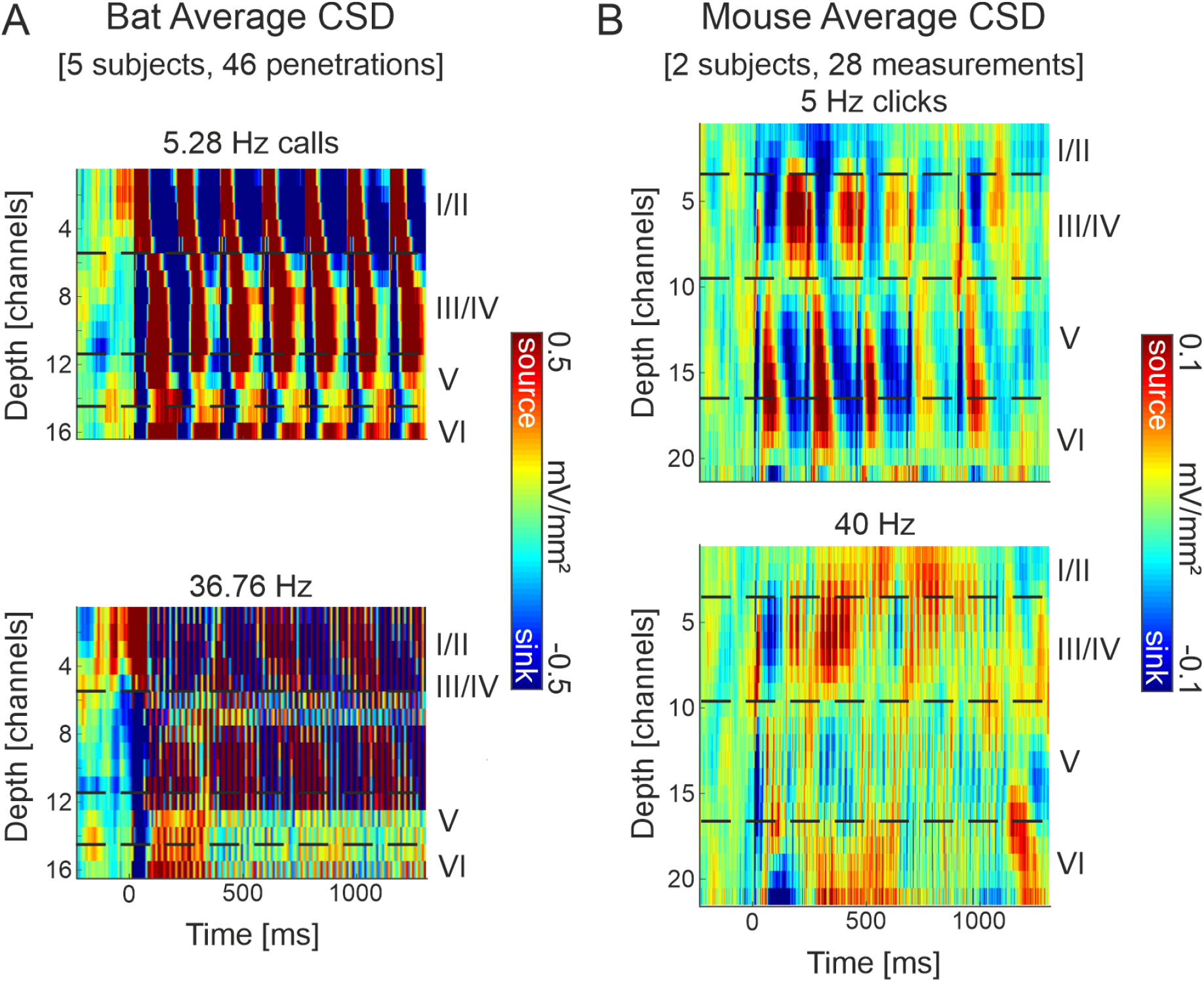
Grand average current-source density profiles. **A:** Seba’s short-tailed bats (n = 46) grand averaged cortical response to a click-like distress call presented repetitively at 5.28 Hz (top) and 36.76 Hz (bottom). **B:** C57/B6 mice (n = 28) grand averaged cortical response to a click train presented at 5 Hz (top) and 40 Hz (bottom). The CSD profiles show the pattern of temporal processing (ms) within the cortical depth (channels are 50 µm apart). Representative layer assignment is indicated with horizontal dashed lines. Current sinks (blue), represent areas of excitatory synaptic population activity, while current sources reflect balancing currents (cf. Happel et al., 2010). Note the different c-axis scales: with much stronger signal from bats, and the different depth scales: slightly thicker cortex for mice, ~20 channels or ~1 mm, than bats 16 channels or ~750 µm.

**Figure 2.**
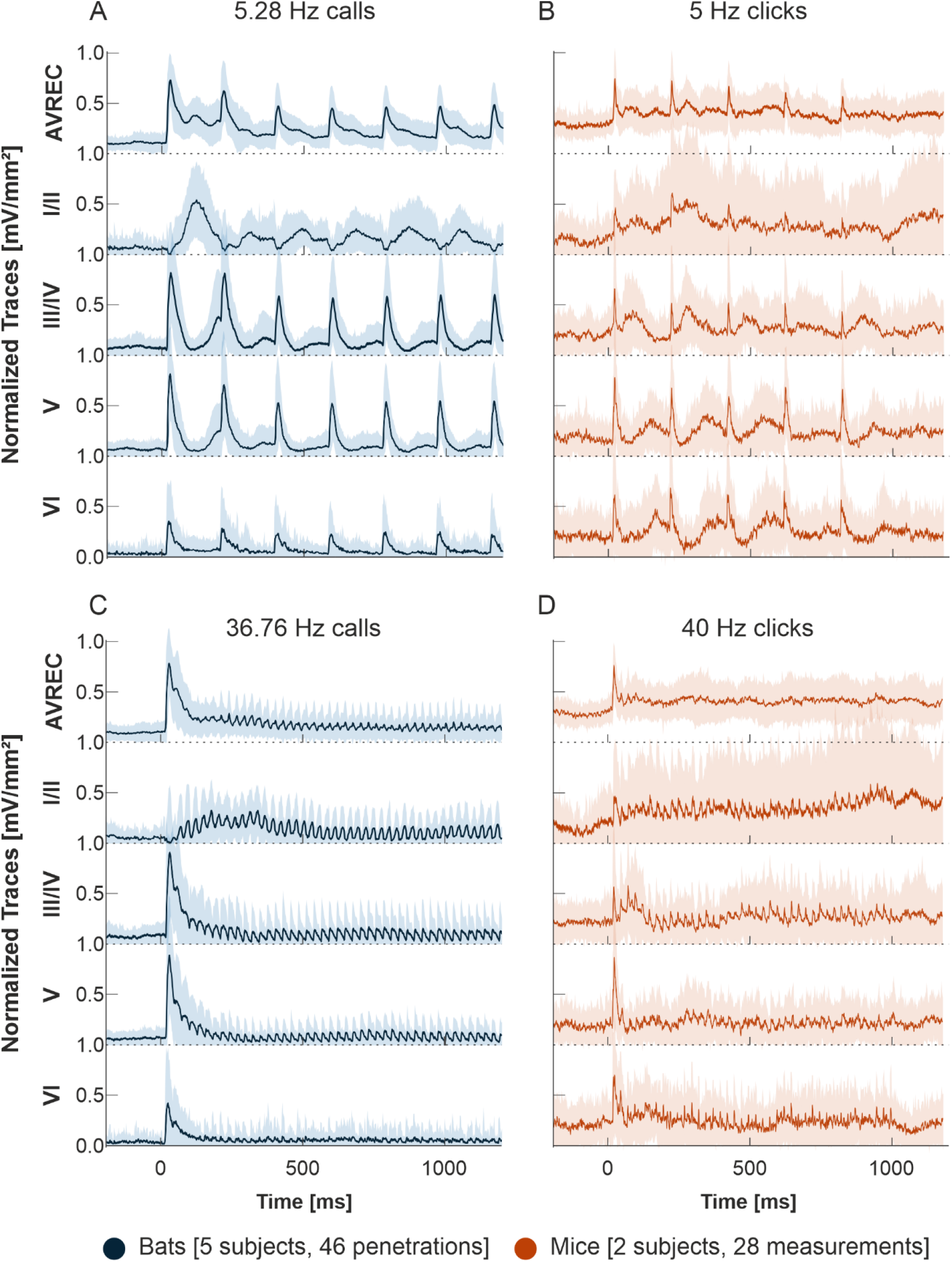
AVREC and layer traces. **A, C:** Bat averaged auditory cortex AVREC trace (top) and all layer traces (I/II, III/IV, V, VI in descending order), in response to **A:** 5.28 Hz or **C:** 36.76 Hz click-like distress calls (blue). **B, D:** Mouse averaged auditory cortex AVREC layer traces, in response to **B:** 5 Hz or **D:** 40 Hz click-trains (orange). Layer traces were calculated on sink activity only (sources replaced with NaN). Confidence intervals are shown in SEM. Traces were all normalized per measurement (bat n=46, mouse n=28). Normalization was done by dividing traces by the first detected peak of the AVREC at 2 Hz (not shown).

### Pharmacology

For mice, pentobarbital (Nembutal, H. Lundbeck A/S, Valby, Denmark) was administered at the onset of surgery with an intraperitoneal injection of 50 mg/kg and supplemented by 20% every hour. Anesthetic depth was regularly checked (every 10-15 min) by paw withdrawal reflex, tail pinch, and breathing frequency. Body temperature was kept stable at 37°C.

Mice received analgesic treatment with Metacam substituted by 5% glucose solution 30 minutes before the end of surgery with 0.3 ml/kg and for 2 days post-operatively with 0.2 ml/kg.

For bats, ketamine-xylazine was administered at surgery onset (ketamine: Ketavet, 10 mg/kg, Pfizer; xylazine: 38 mg/kg). For surgery and for any subsequent handling of the wounds, a local anesthetic (ropivacaine hydrochloride, 2 mg/ml, Fresenius Kabi, Germany) was applied in the scalp.

### Surgery and probe implantation

For mice, the right auditory cortex was exposed by craniotomy and the A1 was located by vascular landmarks. A small hole was drilled on the contralateral hemisphere, over the visual cortex, for implanting a stainless-steel reference wire (Ø 200µm). A recording electrode with a flexible bundle between shaft and connector (A1x332-6mm-50-177_H32_21mm, Neuronexus, Ann Arbor, Michigan USA) was inserted perpendicularly in the A1 and secured with UV-curing glue (Plurafill flow, Pluradent GmbH & Co. KG Magdeburg, Germany). To protect the exposed region of the cortex, the hole was filled with a small drop of an artificial dura replacement compound (Dura-gel, Cambridge Neuro-Tech, Cambridge UK) before being encapsulated. A 3D printed headplate was secured to the top of the exposed skull with dental cement (Paladur, Heraeus Kulzer GmbH, Germany) and the connector (H32-omnetics, Neuronexus) was glued to the top of this headplate with a UV-curing glue and dental cement. Animals were allowed to recover for at least 3 days before habituation to their head-fixation setup.

For bats, whose dataset was previously described in García-Rosales et al. (2020), A1s were exposed by craniotomy (ca. 1 mm^2^) performed with a scalpel blade. A head post (1 cm length, 0.1 cm diameter) was fixed to the top of the skull with dental cement (Paladur). The animals were allowed to recover for at least two days before recording. Per recording session, an acute recording electrode (A1x16-50-177, NeuroNexus), held by micromanipulator, was inserted perpendicularly into the A1 until the uppermost channel was barely visible at the cortical surface.

### Electrophysiological recordings

Mice were placed on a head-fixation treadmill (designed in lab) for 5 days of habituation (from 15 to 75 minutes head-fixed). This treadmill was in a Faraday-shielded acoustic soundproof chamber with a speaker (Tannoy arena satellite KI-8710-32, Tannoy Germany) located 1 m from the head-fixation platform. Recorded local field potentials (LFPs) were fed via an Omnetics connector (HST/32V-G2O LN 5V, 20x gain, Plexon Inc., Dallas, Texas USA) into a PBX2 preamplifier (Plexon Inc.) to be pre-amplified 500-fold and band-pass filtered (0.7-300 Hz). Data were then digitized at a sampling frequency of 1000 Hz with the Multichannel Acquisition Processor (Plexon Inc.). After habituation, mice were head-fixed on the treadmill for 7 consecutive days to record cortical responses to click trains (stimuli duration: 999 ms; click presentation frequency: 5 and 40 Hz; inter-stimulus-interval: 200 and 25 ms respectively; inter-trial-interval: 3 s; carrier tone: pre-determined auditory best frequency; 50 pseudorandomized repetitions; 90 dB sound pressure level; 15 min per measurement) and spontaneous activity (~2 min; no stimuli while recording brain activity from this area). The best frequency, or the tone to which the cortical column responded most strongly based on columnal root mean square, was determined at the beginning of the first day and verified in each consecutive day through tonotopy measurement (pseudo-randomized pure-tones over six octaves: 1 Hz to 32 kHz, tone duration: 200 ms, inter-stimulus interval: 800 ms, 50 pseudorandomized repetitions, 75 dB sound pressure level). Stimuli in this setup were generated in Matlab (Mathworks, R2006b), converted into analog (sampling frequency 1000 Hz, NI PCI-BNC2110, National Instruments), routed through an attenuator (g-PAH Guger Technologies, Graz, Austria), and amplified (Thomas Tech Amp75, Tom-technology, Ilirska Bistrica, Ljubljana). A microphone and conditioning amplifier were used to calibrate acoustic stimuli (G.R.A.S. 26AM and B&K Nexus 2690-A, Brüel&Kjær, Naerum, Denmark). Each subject had 2 click-train measurements per day, totaling 28 for the group, and 2-3 spontaneous measurements per day, totaling 35 for the group (see Table 1).

Bats were placed in a custom-made holder in a Faraday sound-proof chamber and kept at a constant body temperature of 30°C with a heating blanket (Harvard, Homeothermic blanket control unit). A speaker (NeoCD 1.0 Ribbon Tweeter; Fountek Electronics, Hong Kong, China) was positioned 12 cm away from the bat’s right ear. Recordings were made in the left A1. After penetration, per recording session, the probe was connected to a micro preamplifier (MPA 16, Multichannel Systems MCS GmbH, Reutlingen, Germany), connected to an integrated amplifier and analog-to-digital converter with 32-channel capacity (model ME32 System, Multi Channel Systems MCS GmbH). Acoustic stimulation, delivered by Matlab (R2009b), were trains of a single distress syllable (see Figure 1b and d of García-Rosales et al. (2020) for spectrogram; stimuli duration: 2 s; click presentation frequency: 5.28 or 36.76 Hz; inter-stimulus-interval: 189.39 and 27.02 ms respectively; inter-trial-interval: 1 s; 50 pseudorandomized repetitions; intensity: 70 dB SPL rms). Auditory stimuli were digital-to-analog converted using a sound card (M2Tech Hi-face DAC, Pisa Italy, 32 bit; sampling frequency: 192 kHz) and amplified (Rotel power amplifier, model RB-1050, Rotel Europe, West Sussex, England). Spontaneous activity was also recorded at the beginning of each session for 2+ minutes. Bats were recorded from a varying number of times, ranging from 5 to 18 recording sessions. From 5 bats, there was a total of 46 probe penetrations/measurements (see Table 1).

### Current Source Density Analysis

Based on the recorded laminar local field potentials, the second spatial derivative was calculated in Matlab (R2016a-R2022a), yielding the CSD distribution as seen in equation 1:

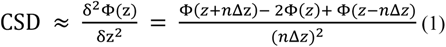

where *ϕ* is the field potential, *z* is the spatial coordinate perpendicular to the cortical laminae, *Δ_z_* is the sampling interval, and *n* is the differential grid (Mitzdorf, 1985). LFP profiles were smoothed with a weighted average (Hamming window) of 9 channels which corresponds to a spatial kernel filter of 450 µm (Happel et al., 2010). CSD distributions reflect the local spatiotemporal current flow of positive ions from extracellular to intracellular space evoked by synaptic populations in laminar neuronal structures. Current sinks thereby correspond to the activity of excitatory synaptic populations, while current sources mainly reflect balancing return currents. Early synaptic thalamocortical inputs persist after intracortical silencing with the GABA_A_-agonist muscimol related to thalamocortical projections on cortical layers III/IV and Vb/VIa (Brunk et al., 2019; Deane et al., 2020; Happel et al., 2010, 2014; Happel & Ohl, 2017) in accordance with reports by others (Schaefer et al., 2015). Early current sinks in the auditory cortex are therefore indicative of thalamic input in granular layers III/IV and infragranular layers Vb/VIa (Happel et al., 2010; Szymanski et al., 2009). Cortical layer designations in mice were made in consideration of *in vitro* (Yamamura et al., 2017) and anesthetized preparations (Deane et al., 2022; Muramatsu et al., 2019), with the difference in anesthetized and awake CSD profiles demonstrated previously in Mongolian gerbils (Deane et al., 2020) for comparison. Layers were selected per penetration in bats and per subject in mice (see Table 1).

CSD profiles were further transformed by averaging the rectified waveforms of each channel by equation 2:

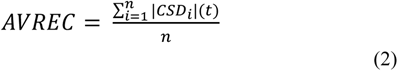

where *n* is the individual channel and *t* is time in ms. This measure gives us the overall temporal local current flow of the columnar activity (Givre et al., 1994; Schroeder et al., 1998). Layer traces were calculated by averaging channel activity in each layer after replacing source activity (positive) with NaN, leaving only sink activity (negative), and then multiplying by −1. After AVREC and layer traces were calculated, they were normalized per measurement. Normalization was done by dividing each AVREC and trace in the measurement by the first detected peak of the AVREC at 2 Hz (not shown) for that measurement. These were then averaged for group images (bat n=46, mouse n=28) and normalized traces were used for peak detection for the following model fit analysis.

All analysis on CSD profiles, AVREC, and layer traces was done at the level of measurements (bat n=46, mouse n=28) rather than subjects (bat n=5, mouse n=2) due to the data collection method in bats: where each measurement was at a new penetration site.

### Model Fit Analysis

A model fit analysis was performed on the averaged peak amplitudes after peak detection on measurement-averaged traces (n=28 for mice, n=46 for bats, see Table 1). Peak detection was calculated with the *max* function in Matlab within detection windows after each stimulus in a presented stimulus train (e.g., for 5 Hz click stimulus, 5 peaks were detected—1 peak after each click). For each of the AVREC and layer trace peak amplitude datasets, 2 models were fitted: exponential decay seen in equation 3 and linear regression seen in equation 4.

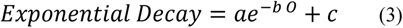

where *a* + *c* is the intercept (the first observed peak amplitude), meaning *a* is depth or the distance between the first observed amplitude and *c*, *b* is the rate of decay (the greater the value, the steeper the decay), *c* is the offset (the value at which the model attenuates), and *O* is the order of peak amplitudes.

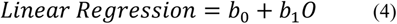

where *b_0_* is the intercept, *b_1_* is the slope, and *O* is the order of peak amplitudes. We used the function *minimize* from the Python SciPy package (Virtanen et al., 2020) to estimate the model parameters. The function used Broyden-Fletcher-Goldfarb-Shanno algorithm to minimize the root mean square error (RMSE), in equation 5:

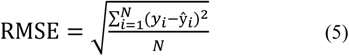

where *y_i_* is the actual value, *ŷ_i_* is the estimated value by the model, and *N* is the number of data points. Model fits and detected peaks were then plotted with overlaid model parameters and RMSE value. Note that indexing in python meant the models started at index 0 but plotting starts at value 1 call or click.

### Phase Amplitude Coupling Analysis

Phase amplitude coupling (PAC) was calculated for each stimulus frequency and on spontaneous activity per measurement (n=28 for mice, n=46 for bats; spontaneous: n=35 for mice, n=46 for bats, see Table 1), based on methodology by Kikuchi et al. (2017) and García-Rosales et al. (2020). LFP signals were filtered in the following low frequency bands with a 4th order bandpass Butterworth filter (Matlab function *filtfilt*): 1 to 3, 3 to 5, … 13 to 15 Hz. LFP signals were also filtered in the following high frequency bands: 25 to 35, 30 to 40, … 95 to 105 Hz. Hilbert transform was applied during the time window of stimulus presentation and, in the stimulus conditions, the average of across trials for the current stimulus and measurement was subtracted from the individual response of each trial to reduce the effect of stimulus-evoked cortical response. Instantaneous phase [φ (t)] for low frequencies and amplitude [A(t)] for high frequencies was then extracted.

To minimize the effect of phase non-uniformities (clustering) in the signal caused by non-oscillatory periodicities in the field potentials, the mean vector of the phase angles was linearly subtracted from the instantaneous phase time series with equation 6:

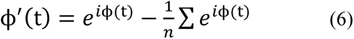

where *ϕ^’(t)* denotes the corrected (de-biased) phase at time t, and n represents the number of series time points. With *ϕ^’(t)* and *A(t)*, a composite time series *z(t)=A(t) × ϕ^’(t)* was constructed. From *z(t)*, the modulation index (MI) was quantified with the following equation 7:

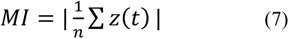

PAC is susceptible to a number of biases in how it is calculated and on the structure of the input signal. A direct comparison between species resulted in very different MI scores across PAC calculations at different frequency pairings. We therefore also computed a surrogate MI by matching the phase series of a given trial with amplitude series of another trial and recalculating surrogate MIs (n = 500) to create a distribution against which we compared observed MI scores (see García-Rosales et al., 2020 Figure 3a). Observed MI scores were z-normalized to the surrogate distribution to obtain the z-scored MI (zMI). If no effect of PAC existed in the data, zMI values would hover around 0, whereas coupling effects would yield zMIs significantly higher than 0 (z-score > 2.5). These zMI values were then arranged into a matrix of high frequency amplitude over low frequency phase PAC pairings and these matrices were used for measurement-normalized comparisons between species over cortical layers. Regions of interest (ROI) were determined based on each species strongest area of PAC, and a clustermass permutation analysis was run on both ROIs per comparison (see below).

**Figure 3.**
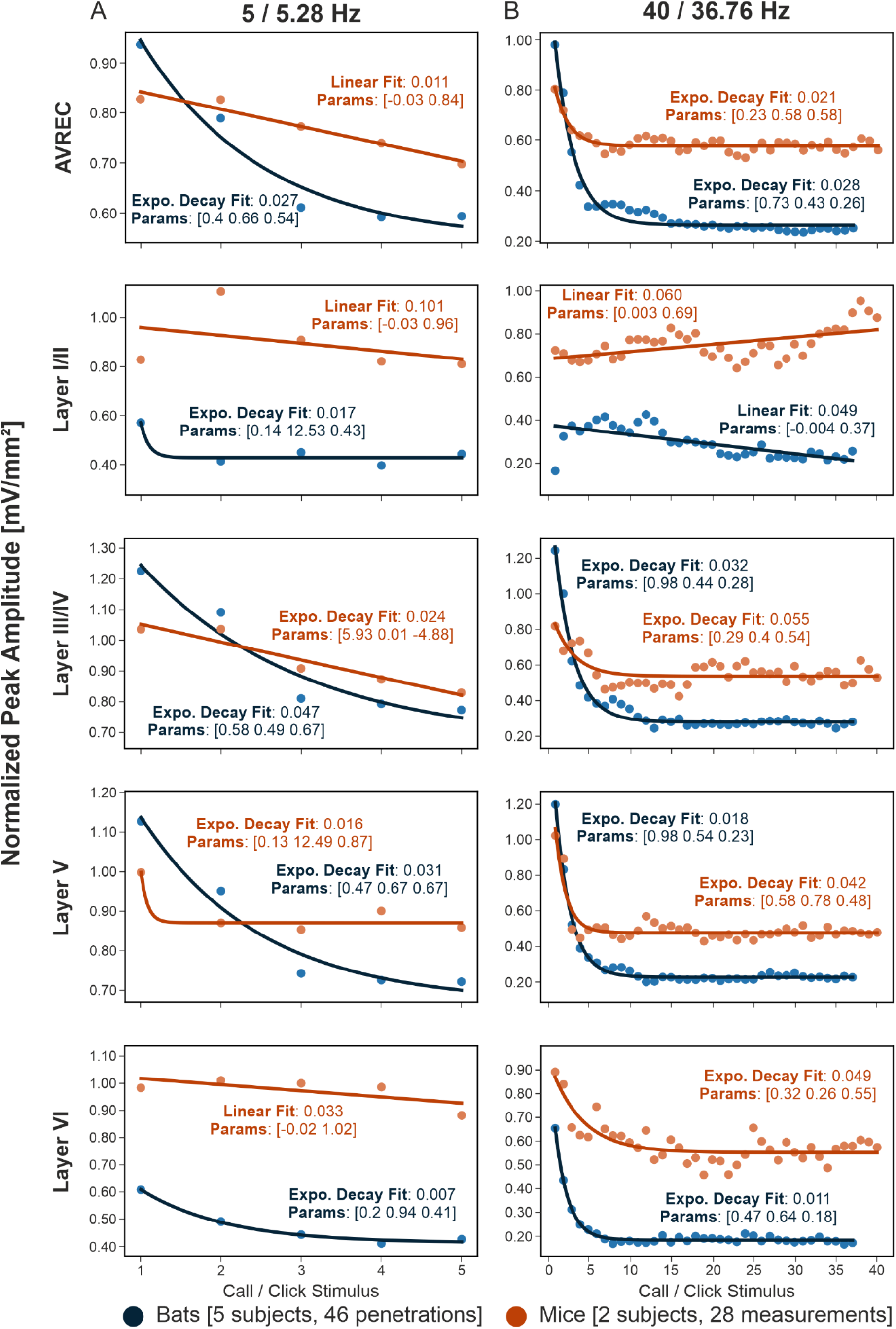
Model Fit Analysis. **A:** Bat (blue, n=46) and mouse (orange, n=28) averaged response peak amplitudes over consecutive stimulus repetition of 5 or 5.28 Hz with overlaid model fit. **B:** Bat and mouse group-averaged response peak amplitude over consecutive stimulus repetition of 40 or 36.76 Hz with overlaid model fit. The model selected, exponential or linear decay is overlaid as a trace, along with the fit value calculated by RMSE and the model parameters. The closer to zero that the model fit is, the better. For expo.: parameters are [dynamic range, rate of decay, offset]. For linear: [slope, intercept]

### Clustermass Permutation Analysis

The clustermass permutation analysis is specifically suited to control for a familywise error rate (FWER; cf. Groppe et al., 2011). To perform this analysis on the level of measurement (n=28 for mice, n=46 for bats, see Table 1), we extracted a *t* statistic pointwise across matrices in comparison of groups and pre-selected a significance *t* threshold based on a two-tailed p-value < 0.05. Any statistic result at or above this significance threshold was converted to a 1 and anything below was converted to a 0—creating a binary matrix of 0s and 1s, where 1 is a possible point of significance in the comparison of those matrices. The 1s within each ROI were then summed to create our observed clustermass values. Next, we permuted the groups 500 times; condition containers were created, equal to observed group sizes, and the measurements from both groups were randomly allocated into those containers. The same point-wise statistic-and-threshold-calculated binary map was produced for each permutation with the total sum of 1s for each ROI taken as a permutation clustermass value. This created a distribution of 500 permutation clustermass values to which the observed clustermass could be compared. A p-value was calculated according to where the observed clustermass value fell onto the permutation distribution. This test indicates if the difference in the observed conditions is significant above chance, if p < 0.05—or put another way, it tells us how reliable the observed results are.

#### Continuous Wavelet Transform Analysis

Spectral analysis was performed in Matlab using the wavelet analysis toolbox function *CWT* (short for Continuous Wavelet Transform) for the following variables: animal, condition, stimulus, and recorded signal. Important parameters fed into the CWT were as follows: layer channels from CSD profiles, frequency limits: 5 to 100 Hz (below the Nyquist), and wavelet used: analytic Morse (Lilly & Olhede, 2012; Olhede & Walden, 2002). For layer-wise wavelet analysis, the center channel of each layer was fed into the CWT. A trial-averaged scalogram was calculated for each cortical layer and wavelet power—per frequency, per time point—for each measurement (n=28 for mice, n=46 for bats, see Table 1) with equation 8.

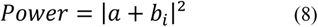

where *a + b_i_* represents the complex number output of the trial-averaged CWT analysis (Lachaux et al., 1999). Power was normalized to the maximum power in each measurement to result in a relative power of signal to background noise and to account for the large species difference in scale (bats had stronger unnormalized power by a factor of 3, not shown). Single trial scalograms were calculated for each measurement as well and, on these, phase coherence—per frequency, per time point—for each subject was computed with equation 9:

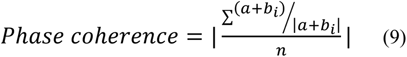

Normalized power and phase coherence data were averaged pointwise across measurements for group plots. Clustermass permutations (described above) were performed for the difference between spectral representations in each layer, with an ROI of 200 ms of baseline activity the point of onset of the second stimulus (e.g., 5 /5.28 Hz: −200 to 189 ms) in each frequency band. Frequency bands were split as follows: theta 4-7 Hz, alpha 8-12 Hz, low beta 13-18 Hz, high beta 19-30 Hz, low gamma 31-60 Hz, and high gamma 61-100 Hz. For power calculations: the test statistic for permutation was the student’s t test and a Cohen’s D matrix was generated to indicate effect size per frequency at each time point. For phase coherence calculations: the test statistic for permutation was the non-parametric *mwu* (Cardillo, 2009; Maris et al., 2007) test and effect size, r, was indicated with the z score output as in equation 10:

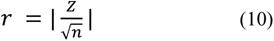

## Results

### Shared microcircuitry but differing cortical response profiles

Seba’s short-tailed bats and C57/B6 mice have a similarly thick auditory cortex, but it is slightly thicker in mice (~1 mm; Chang & Kawai, 2018) than in bats (~750 µm; García-Rosales et al., 2019). Figure 1 shows the group averaged CSD profiles for bats and mice at ~5 and ~40 Hz stimulus presentation (see Supp Figure 4 for averaged LFP profiles and Supp Figure 5 for an anatomical comparison of the averaged group CSDs to the above references). Awake, head-fixed bats heard a species-specific distress syllable repeated at 5.28 and 36.76 Hz, over 2 seconds (1 second of stimulus presentation is shown here for comparison; see García-Rosales et al. 2020). Awake, head-fixed, freely moving mice were presented with broadband click trains at 5 and 40 Hz over 1 second.

**Figure 4.**
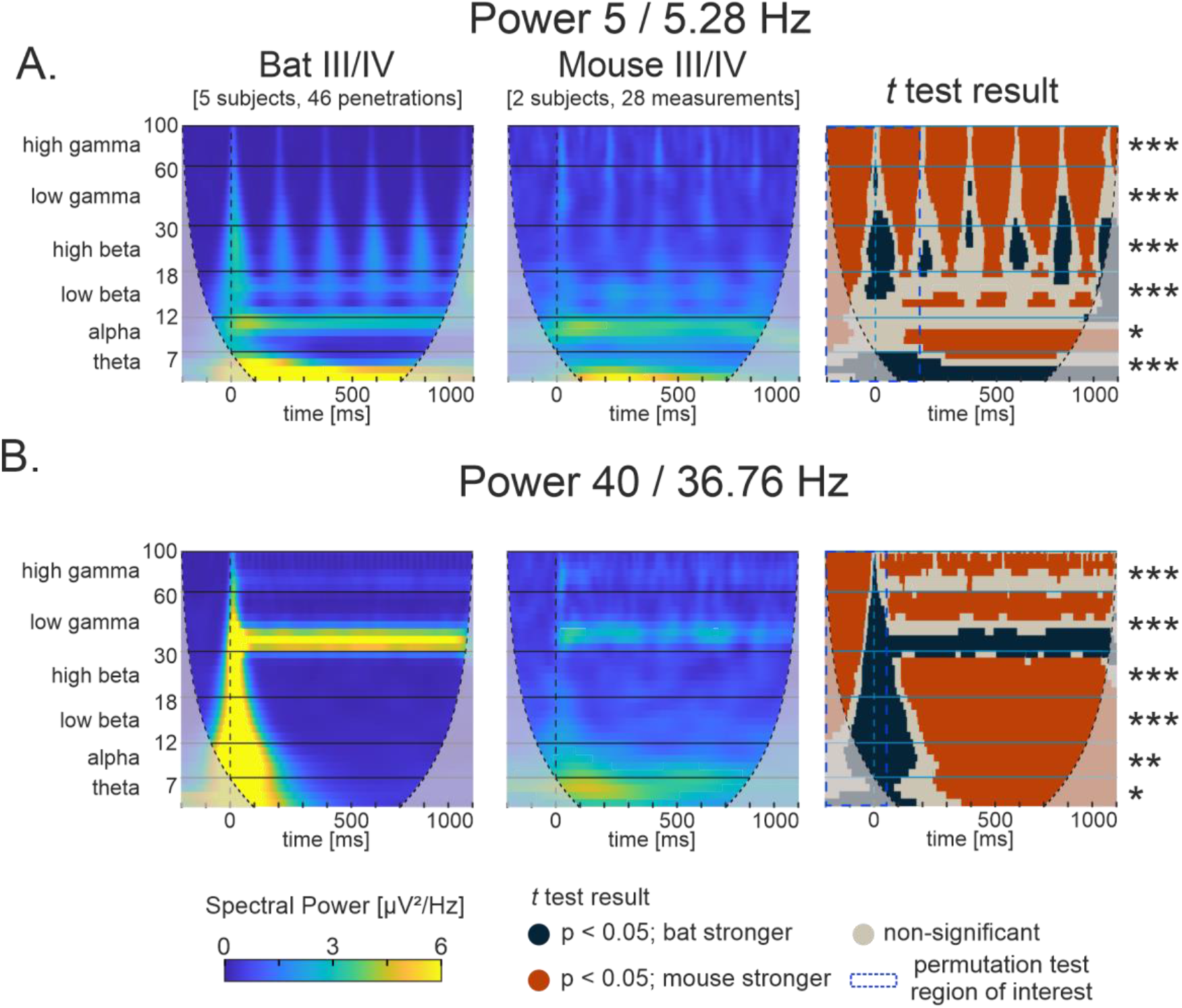
Normalized power scalograms of continuous wavelet transform. **A, B:** bat (left, n=46) and mouse (middle, n=28) average normalized power CWT profiles of layer III/IV during **A:** ~5 Hz and **B:** ~40 Hz stimuli presentation. Power was normalized to the maximum power per measurement. The two-sided Student’s *t* test result plots (right; df: 72) in comparison between bat and mouse normalized power scalograms show contrast between where bats have significantly higher normalized spectral power (blue, p<0.05) and where mice have significantly higher normalized spectral power (orange, p<0.05). ROIs (dashed blue boxes) are overlaid where clustermass permutation analysis was run in each oscillatory band (**A:** from −200 to 189 ms and **B:** from −200 to 25 ms; where 0 ms is stim onset). Horizontal borders designated spectral frequency bins: theta: 4-7 Hz (skipping delta in this analysis), alpha: 8:12 Hz, beta low: 13:18 Hz, beta high: 19:30 Hz, gamma low: 31:60, gamma high: 61:30. Permutation results are shown as significance stars to the left, indicating results as reliable with their respective ROI (*=p<0.05, **=p<0.01, ***=p<0.001). See Table 2 for corresponding *p* values.

**Figure 5.**
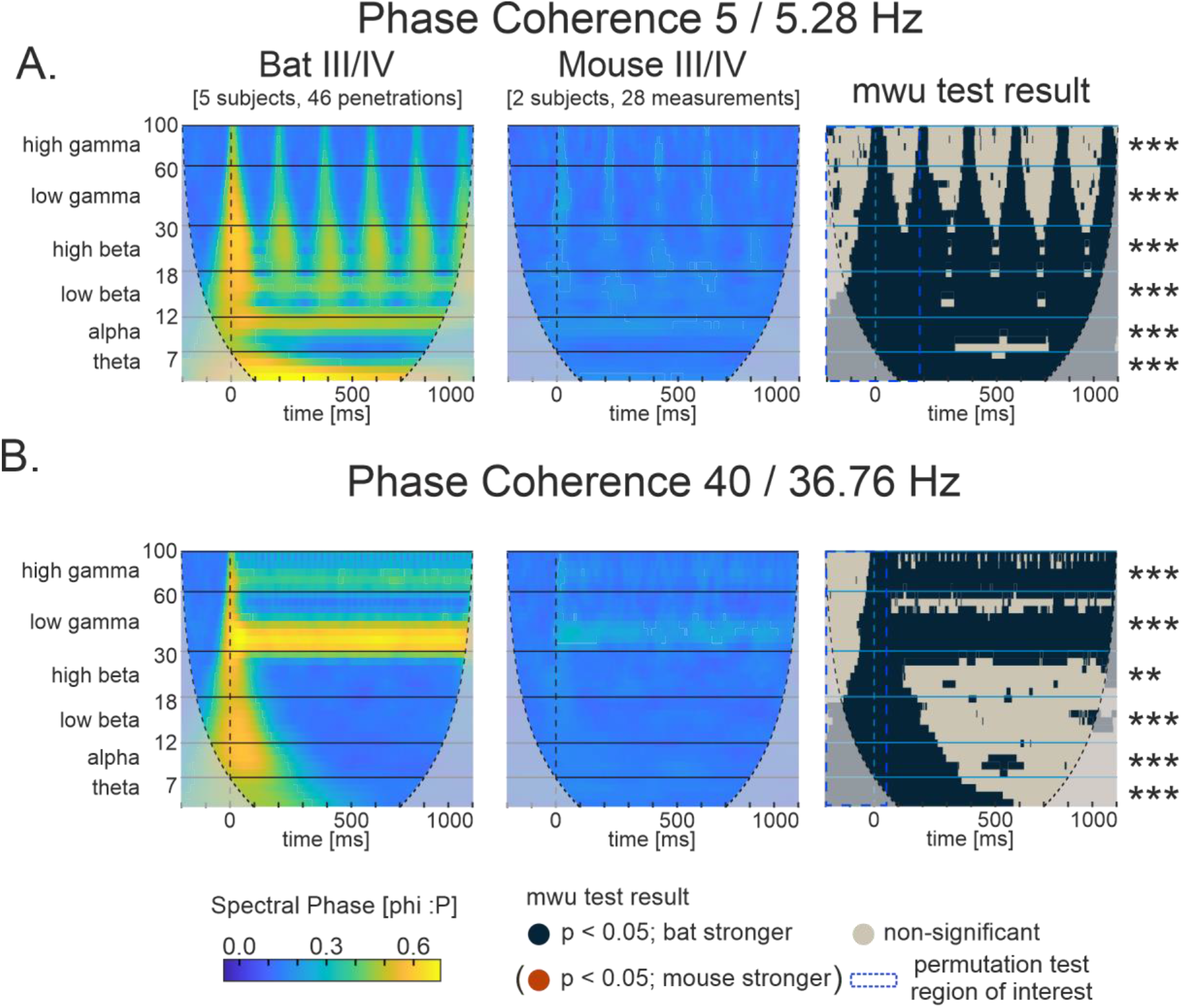
Phase coherence scalograms of continuous wavelet transform. **A, B:** bat (left, n=46) and mouse (middle, n=28) average phase coherence CWT profiles of layer III/IV during **A:** ~5 Hz and **B:** ~40 Hz stimuli presentation. The two-sided *mwu* test result plots (right, df: 72) in comparison between bat and mouse phase coherence scalograms show contrast between where bats have significantly higher normalized spectral phase coherence (blue, p<0.05) and would have shown where mice had significantly higher normalized phase coherence (orange, p<0.05, in this case not present). ROIs (dashed blue boxes) are overlaid where clustermass permutation analysis was run in each oscillatory band (A: from −200 to 189 ms and B: from −200 to 25 ms; where 0 ms is stim onset). Horizontal borders designated spectral frequency bins: theta: 4-7 Hz (skipping delta in this analysis), alpha: 8:12 Hz, beta low: 13:18 Hz, beta high: 19:30 Hz, gamma low: 31:60, gamma high: 61:30. Permutation results are shown as significance stars to the left, indicating results as reliable with their respective ROI (*=p<0.05, **=p<0.01, ***=p<0.001). See Table 2 for corresponding *p* values.

**Table 2.** Between group CWT spectral band clustermass comparison. Corresponding to **Figure 4** &**Figure 5**. Bat (n=46) vs Mouse (n=28) ROI comparisons of power and phase coherence scalograms for ~5 and ~40 Hz. ROI was the 200 ms before stimulus onset to 189 ms (for 5.28 Hz) or 25 ms (for 40 Hz) after stimulus onset in this condition for each spectral band. The test was a two-sided Student’s *t* test on spectral power values, and a two-sided *mwu* test on phase coherence values. Each test had degrees of freedom: 72. *p* value results and corresponding mean and std are shown. In bold are *p* values where significant above chance (p < 0.05).

|  |  | Normalized Power |  |  | Phase Coherence |  |  |
| --- | --- | --- | --- | --- | --- | --- | --- |
|  |  | *p* value | mean | std | *p* value | mean | std |
| $\sim 5$ Hz | theta | **0.000** | 116.776 | 201.918 | **0.000** | 110.574 | 369.175 |
|  | alpha | **0.028** | 94.676 | 176.134 | **0.000** | 94.714 | 265.334 |
|  | beta low | **0.000** | 83.174 | 127.432 | **0.000** | 89.876 | 211.257 |
|  | beta high | **0.000** | 86.382 | 134.406 | **0.000** | 87.820 | 181.234 |
|  | gamma low | **0.000** | 157.592 | 212.520 | **0.000** | 145.158 | 166.716 |
|  | gamma high | **0.000** | 130.882 | 205.087 | **0.000** | 125.694 | 71.814 |
| $\sim 40$ Hz | theta | **0.014** | 72.490 | 162.299 | **0.000** | 62.692 | 186.233 |
|  | alpha | **0.008** | 53.774 | 125.424 | **0.000** | 40.466 | 131.779 |
|  | beta low | **0.000** | 58.204 | 116.705 | **0.000** | 47.138 | 102.103 |
|  | beta high | **0.000** | 50.516 | 86.014 | **0.002** | 52.118 | 86.863 |
|  | gamma low | **0.000** | 88.592 | 113.666 | **0.000** | 85.342 | 86.573 |
|  | gamma high | **0.000** | 69.468 | 96.232 | **0.000** | 74.210 | 51.314 |

The supragranular layer of the bat A1 was proportionally much thicker than that found across mice and had a very strong and consistent following response which lagged behind the thalamocortical response (Figure 1A; Supp Figure 5). In the mouse average CSD profile (Figure 1B), the granular sink was very light in comparison to the early infragranular response. Where we saw very clear following responses down the depth of the cortex in bats at a lower (5.28 Hz) and higher (36.76 Hz) frequencies, the following response in mice was noisier—less distinct sink and source areas standing out of the background activity—and more relegated to thalamic input areas, with separate, repeated granular and infragranular sinks following the stimuli. The noisier signal seen in awake mice was not surprising in comparison with the classically less noisy ketamine-anesthetized signals in mice as well as Mongolian gerbils (Deane et al., 2020, 2022), due to ketamine being a neuronal synchronizer. Interestingly, the awake bat cortical activity was then less noisy compared to these awake rodent datasets (looking more similar to the anesthesia-induced synchrony in data cited above), in the sense of legible sinks far above the baseline of cortical activity for each stimulus presentation.

### Accurate cortical following responses through all layers in bats

Figure 2 shows the averaged Average Rectified CSD (AVREC) and layer traces per group for ~5 and ~40 Hz. Amplitude was normalized to each measurement’s first AVREC peak detected in their 2 Hz conditions. This visually represents the relative contribution of the layers to the full cortical column activity. Single animal, non-normalized traces can be found in Supp Figure 1 Supp Figure 2 for bats and Supp Figure 3 for mice.

Sinks originating in the granular layer of mice often spread up into supragranular layers, causing a stimulus-locked, small amplitude supragranular response at tone onset (Figure 1). In the bats, the supragranular layer consistently lagged the stimulus-locked thalamic input activity of layers III/IV, V, and VI, creating an accurate, though delayed, following response to low and high frequency stimulus presentations. In both bat and mouse ~5 Hz AVREC traces, there was an initial onset response in the full column and a second, smaller and broader peak after both the first and second stimulus responses. In the bats, that second peak was driven almost exclusively by the supragranular activity. In the mice, a second, broader peak was seen in all layers, demonstrating less laminar specificity.

Mouse cortical activity was noisier than bat cortical activity, as noted in CSD profiles. That is to say, while bat averaged traces revealed almost uniform following responses at high and low stimulus presentation frequencies, mouse cortical activity seemed to contain more high frequency jitter, higher levels of background noise, and more variable following response profiles across consecutive stimuli presentation. We quantitatively analyzed this phenomenon in both the model fit and continuous wavelet analyses described below. Qualitatively in Figure 3, it appeared that normalized activity in bats began closer to relative 0 than in mice and then sank back nearer to 0 in each layer after the stimulus response. The only exception was in the first 500 ms of the supragranular activity where there was a distinct slow wave response in bats, and in the AVREC trace which included both sink and source activity rectified.

Because we could see the responses to consecutive stimuli riding the first onset response in the bats at 36.76 Hz, we separated these two components with bandpass filters (Supp Figure 6). We filtered +/- 3 Hz around the stimulus frequency to reveal, more strictly, the following response components and we filtered from 1 to 4 Hz to reveal the onset response. With the stimulus frequency filter, bats layer I/II showed the same response lag and then a consistent amplitude response to consecutive stimuli. In the other layers, there was a relatively consistent higher amplitude following response in the first ~100 ms of tone presentation which then attenuated to a more even response to consecutive stimuli. In mice at 40 Hz, stimulus response and attenuation to consecutive responses was also more variable in mice when filtered around stimulus presentation frequency. With the onset component filter, the onset in bats was consistently higher than mice in the AVREC, granular, and infragranular layer traces. The supragranular layer was nearly flat in bats but the previously described, lagged, slow wave was visible in both species with this filter.

**Figure 6.**
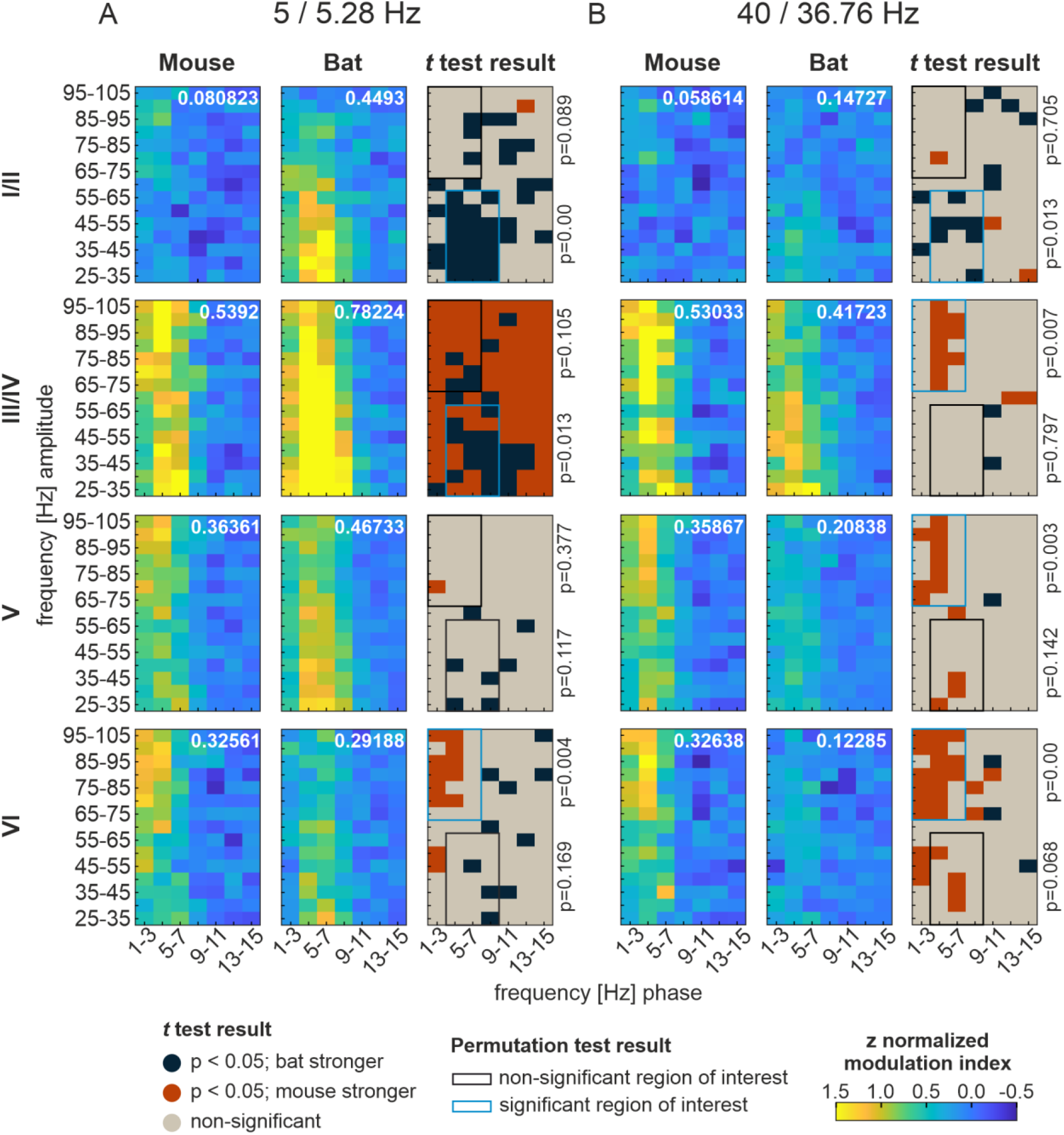
Phase amplitude coupling profiles over low and high frequency. The PAC profiles, high frequency amplitude over low frequency phase, of mice (left, n=28) and bats (middle, n=46) for each cortical layer center channel (I/II, III/IV, V, VI from top to bottom) for **A:** ~5 Hz and **B:** ~40 Hz are represented as the z-score normalized modulation (zMI) index. Higher zMI indicates better coupling. The Student’s *t* test result plots (right, df: 72) in comparison between bat and mouse PAC profiles show contrast between where bats have significantly higher zMI (blue, p<0.05) and where mice have significantly higher zMI (orange, p<0.05). Overlaid are ROI boxes where permutation clustermass analysis was calculated, blue if significant (p<0.05), black if nonsignificant, with corresponding p values on the right, indicating reliability of *t* tests within these areas. See Table 3 for corresponding *p* values.

**Table 3.** Between group PAC ROI comparison. Bat (n=46) vs Mouse (n=28) PAC profiles at delta-high gamma (1-7 Hz phase vs 65-105 Hz amp) and theta-low gamma (3-9 Hz phase vs 25-65 Hz amp) phase-amp couplings. The test was a two-sided Student’s *t* test on output zMI, corresponding to **Figure 6** & **Figure 7**. Regions were chosen based on the PAC profiles and not exact spectral frequency bins. Comparisons were done with the same regions across all layers and stimulus conditions. Each stimulus presentation test had degrees of freedom: 72, and each spontaneous test had degrees of freedom: 79. *p* value results and corresponding mean and std are shown. In bold are *p* values where significant above chance (p < 0.05).

| Frequency | Layer | Region of Interest | PAC Permutation |  |  |
| --- | --- | --- | --- | --- | --- |
|  |  |  | *p* value | mean | std |
| 5 / 5.28 Hz | II | Delta / high gamma | 0.089 | 2.548 | 2.057 |
|  |  | Theta / low gamma | **0.000** | 2.300 | 2.925 |
|  | III/IV | Delta / high gamma | 0.105 | 2.350 | 2.438 |
|  |  | Theta / low gamma | **0.013** | 2.154 | 2.844 |
|  | V | Delta / high gamma | 0.377 | 2.327 | 2.213 |
|  |  | Theta / low gamma | 0.117 | 2.061 | 2.194 |
|  | VI | Delta / high gamma | **0.004** | 2.413 | 2.047 |
|  |  | Theta / low gamma | 0.169 | 2.070 | 1.989 |
| 40 / 36.75 Hz | II | Delta / high gamma | 0.705 | 2.485 | 1.563 |
|  |  | Theta / low gamma | **0.013** | 2.094 | 1.738 |
|  | III/IV | Delta / high gamma | **0.007** | 2.213 | 2.107 |
|  |  | Theta / low gamma | 0.797 | 2.106 | 2.094 |
|  | V | Delta / high gamma | **0.003** | 2.401 | 1.790 |
|  |  | Theta / low gamma | 0.142 | 1.951 | 1.544 |
|  | VI | Delta / high gamma | **0.000** | 2.390 | 1.843 |
|  |  | Theta / low gamma | 0.068 | 2.110 | 1.444 |
| ~ | II | Delta / high gamma | **0.001** | 2.445 | 2.766 |
|  |  | Theta / low gamma | **0.000** | 2.090 | 3.596 |
|  | III/III/IV | Delta / high gamma | **0.000** | 2.273 | 2.848 |
|  |  | Theta / low gamma | **0.000** | 2.273 | 3.671 |
|  | V | Delta / high gamma | **0.001** | 2.344 | 3.144 |
|  |  | Theta / low gamma | **0.000** | 2.082 | 3.311 |
|  | VI | Delta / high gamma | 0.376 | 2.364 | 2.201 |
|  |  | Theta / low gamma | **0.004** | 2.042 | 2.260 |

### A greater dynamic range in response amplitude to consecutive stimuli in bats

Due to an evident difference in the response profile across consecutive stimuli, including lower background activity compared to signal response and more uniformity in bat responses, we performed a model fit analysis, with 2 models to choose from algorithmically: exponential or linear. Figure 3 shows the averaged peak amplitudes of responses after stimuli overlaid with the model selected and its fit value (root mean square error, RMSE) and parameters. The best fitting model was typically exponential decay. For bats, at both presentation frequencies, exponential decay was selected in all traces except the supragranular layer at 36.76 Hz, where a linear fit was selected. At 5 Hz in mice, layer III/IV and V were the only traces selected for exponential decay. In these layers, the exponential fit was a better choice for the bat dataset: the offset of 5 Hz mouse layer III/IV was well below a possible peak amplitude due to how shallow the rate of decay was, and the rate of decay in 5 Hz layer V was severely steep.

The bat 5.28 Hz III/IV and V models were comparable to each other and spanned a greater dynamic range (intercept – offset) than mouse 5 Hz V. In ~40 Hz, datapoints were more aligned with an exponential fit for bats in every case except the AVREC peak amplitudes. Importantly, bats had a greater dynamic range parameter in the AVREC and layers III/IV through VI, indicating again a consistently deeper suppression of consecutive responses at this higher frequency presentation.

To exclude that the onset component was the main effector for the model fitting, we ran the same analysis with bandpass-filtered signals (Supp Figure 7). The model fits between both species were more similar but they did not explain the data as well as when the onset response component was included for either species. However, the bat data still generally showed a greater dynamic range than mice throughout the layers.

**Figure 7.**
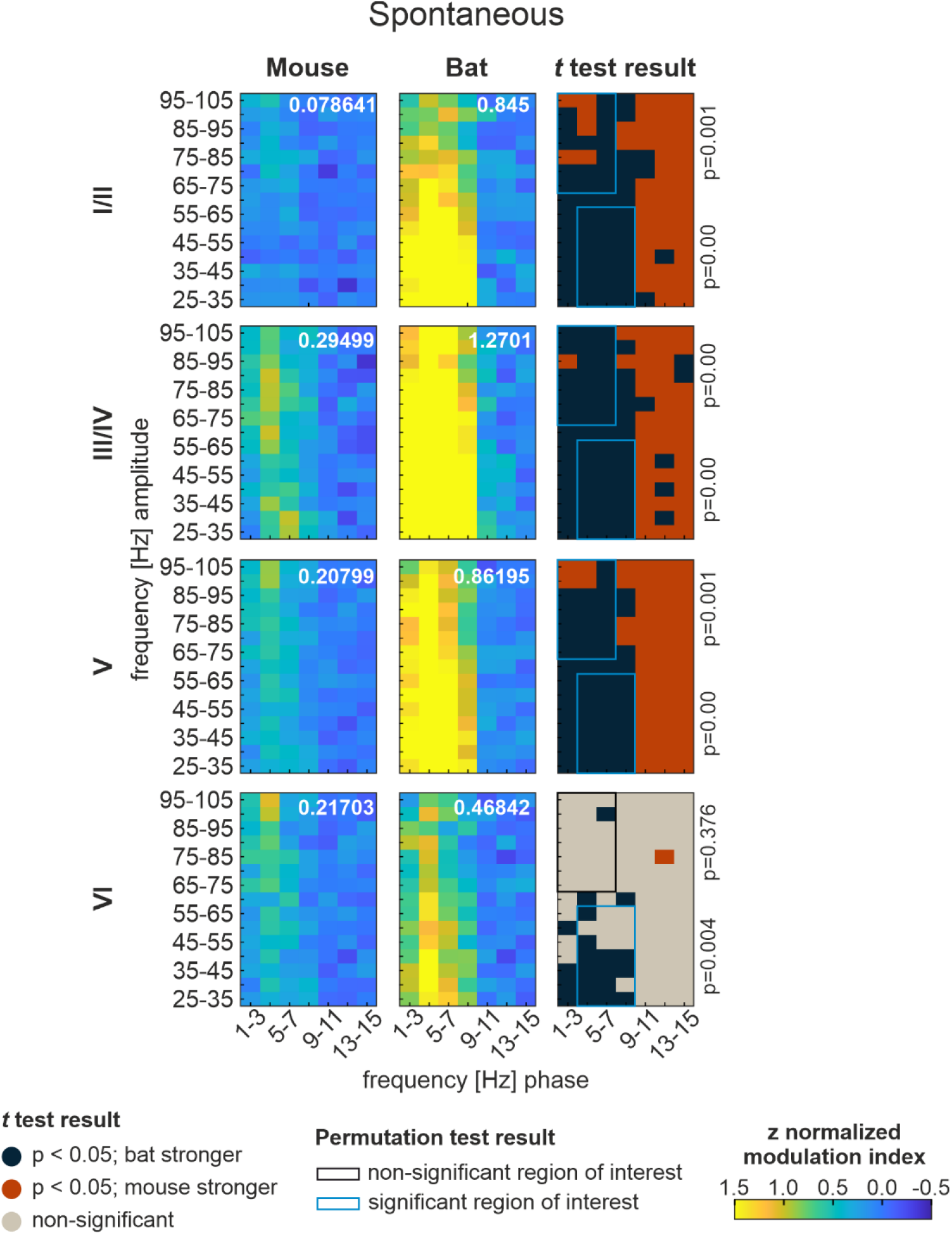
Phase amplitude coupling profiles from spontaneous activity. The PAC profiles, high frequency amplitude over low frequency phase, of mice (left, n=35) and bats (middle, n=46) for each cortical layer center channel (I/II, III/IV, V, VI from top to bottom) for **A:** ~5 Hz and **B:** ~40 Hz are represented as the z-score normalized modulation (zMI) index. Higher zMI indicates better coupling. The Student’s *t* test result plots (right, df: 79) in comparison between bat and mouse PAC profiles show contrast between where bats have significantly higher zMI (blue, p<0.05) and where mice have significantly higher zMI (orange, p<0.05). Overlaid are ROI boxes where permutation clustermass analysis was calculated, blue if significant (p<0.05), black if nonsignificant, with corresponding p values on the right, indicating reliability of t tests within these areas. See Table 3 for corresponding *p* values.

### Better signal to noise ratio in spectral power scalograms and stronger phase coherence in bats

We performed CWT analysis to discern both inter-trial phase variability through the lens of phase coherence and signal to noise ratio through normalized spectral power. Across both normalized power (Figure 4) and phase coherence (Figure 5) for both stimulus conditions and across all spectral frequency bands, observed clustermass was significantly higher than chance according to permutation analysis, attesting to reliability in the observed results.

In comparisons of normalized spectral power (Figure 4), the background around stimulus response was significantly different, with mice showing a higher level of spectral noise relative to signal response. The time at which thalamic input reached the cortex (~15 ms), bats had significantly higher power for each signal response in the beta and low gamma range in the ~5 Hz condition and for the onset signal response in the theta through low gamma range in the ~40 Hz condition. A band of non-significant test results surrounded the higher spectral power of signal response in bats to transition to the higher spectral power of the background in mice. Effect sizes were taken and correspond with level of significance (not shown).

Phase coherence (Figure 5) was significantly stronger in bats across a broadband of spectral frequencies following each stimulus presentation in the ~5 Hz condition and at the onset signal response in the ~40 Hz condition. In this higher frequency condition, there was also significant phase coherence around the stimulus presentation in the low gamma range which was significantly stronger in bats.

### Fundamentally different local and global phase amplitude coupling profiles

We then performed PAC analysis (Figure 6) within cortical layers of both species to differentiate coupling profiles. PAC was strongest in mice centered around delta and high gamma pairings when and strongest in bats around theta/alpha and low gamma pairings. These were chosen as ROIs for comparison between PAC profiles. Due to spatial distinction in the CSD signal, laminar differences were found in both species, with lower or essentially no coupling in supragranular layers and higher coupling in the granular layer. Mouse layer III/IV through VI profiles revealed theta gamma (low and high) coupling at roughly equivalent levels in both high and low stimulus frequency conditions. The bat PAC was stronger during the 5.28 Hz condition compared to the 36.76 Hz condition, possibly indicating a stimulus dependance in bats. However, Bat PAC was only stronger than mouse PAC in layer III/IV in the ~5 Hz stimulus presentation. Generally, mouse PAC in the delta high gamma pairing was significantly higher after clustermass permutation testing.

PAC analysis was performed also on spontaneous, or resting state, activity (Figure 7). During spontaneous activity, bats had PAC at a wider range of low frequency phases and high frequency amplitudes than in stimulus conditions. Bats had significantly higher PAC in both ROIs here.

## Discussion

We compared the cortical response profiles across all layers of A1 in two small mammalian species, one a flying auditory specialist and one a terrestrial olfactory specialist. The nervous systems of both species have been adapted to fill specific ecological niches. Seba’s short-tailed bats have sophisticated social communication and echolocate for navigation (Beetz et al., 2017; García-Rosales et al., 2022; Hechavarría et al., 2013; López-Jury et al., 2021; Thies et al., 1998; Weineck et al., 2020), meaning that temporally accurate perception of auditory signals is paramount to successful social and flying behavior. Mice have a smaller repertoire of social vocal cues (Fonseca et al., 2021) and rely on their whiskers and olfaction for navigation (Gire et al., 2016), indicating that comparably precise cortical representation of auditory cues is not necessary for behavioral success. Through this novel comparison of CSD profiles, we have shown that, in these conditions, Seba’s short-tailed bats have a better signal to noise ratio in auditory response to repetitive stimuli, unique PAC profiles, and less inter-trial phase variability than black 6 mice. We posit that these phenomena are based on a species-specific overall recruitment of the evolutionarily conserved cortical microcircuitry.

### Methodological Considerations

Many model organisms are investigated in neuroscience and interpretation of results must consider necessary abstractions based on species, study design, and methods. One objective of this study was to close the inherent gap between datasets using a comparative data analytical approach. One difference in these datasets was the hemisphere of recording. While data was recorded in the left A1 in bats, we used data from the right A1 in mice. While the lateralized functions of the A1 have been established in many studies (c.f. Wetzel et al., 2008; Zatorre & Belin, 2001), others found no difference between adaptation lifetime between auditory cortex hemispheres (Lu et al., 1993). In mice, a recent study recorded spike trains from the left and right AC and found longer integration time windows in superficial layers of the right AC linked with stronger recurrent excitation (Neophytou et al., 2022). On the other hand, in mustached bats, spike trains were recorded from the left and right hemisphere doppler shifted areas of the A1 and it was found that the left hemisphere had longer integration windows (Kanwal, 2012), contrary to findings in mice. This study therefore demonstrated that the left A1 was better in discriminating call types and that the right was better for temporal processing of echolocation. If this holds for non-doppler shifted A1 areas across bat species, then we would have recorded from each species’ respective hemisphere for longer temporal integration windows and therefore superior spectral integration. However, as a cross-regional and species comparison was beyond the scope of this study, these dependencies should be further explored in future comparisons of this kind. It may be suitable to continue comparative studies in consideration of lateralized function (longer integration windows) instead of strict left or right-side comparisons.

Despite the shared hemispheric “call discrimination” function in respective datasets, the difference in stimuli may partially account for differential recruitment of cortical microcircuitry between species. We used a broadband click for mice and a distress syllable for bats. It has been shown that behavioral relevance and valence distinctly affects processing in the A1 (De Franceschi & Barkat, 2021; Zempeltzi et al., 2020) and further studies along this line collecting new data, rather than comparing pre-collected sets, ought to consider emotional valence when generating stimuli. The challenge of finding comparable stimulus categories that cater to emotional aspects for different species still persists. Given the complexity of ecological consideration, we do not believe the stimuli type used in the two data sets fully account for the described discrepancies, particularly in a study design focusing purely on passive listening (Kato et al., 2015).

### Shared mammalian laminar structure

While the laminar structure of the neocortex is shared between mammals (Mountcastle, 1997), there were some differences between Seba’s short-tailed bat and black 6 mouse primary auditory cortex layers. The supragranular layer of activity revealed by CSD analysis (Figure 1) in bats was consistently thicker—taking up more channels on the probe relative to the full profile—than in mice. The distinction between supragranular and granular activity was clearer in the bat CSD profiles, where the former had a definite lag behind the latter. In histological studies of each’s laminar profiles, layer I was proportionally thicker in the bat A1 (García-Rosales et al., 2019) than in mice (Chang & Kawai, 2018). While this agrees with the population activity we observed in this study, it did not fully account for how much deeper the uppermost sink activity protruded in the cortical depth. This may indicate differing recruitment of layer II neurons to either assist in layer I cross-columnar activity or in the layer III and IV excitation feedback circuitry between species. In the thinner bat A1 cortex, thalamic input to granular and infragranular layers appeared as more of a single input sink. In the mouse A1, there were separable input sinks at the onset of a stimulus and both layer V and VI were thicker than in the bat CSD profiles. We found confirmation of these layer designations again from Chang & Kawai, (2018) and García-Rosales et al., (2019). Therefore, despite shared architecture, there were differences in the proportional layer sizes which likely contributed to the differing recruitment profiles across species. What might also account for the

A possible reason for the overall strength difference is neuronal geometry and cortical architecture. LFP signal is generally stronger the smaller the cortex, because pyramidal cell somas line up into fewer rows in smaller cortices and smaller brains tend to have smaller cell bodies that are more densely packed (Herculano-Houzel et al., 2007). Both of these small brain adaptations vertically align pyramidal neurons for stronger open field dipoles that can be measured as LFP (Buzsáki et al., 2012). This study did not analyze differences in pure strength of activity. However, the stronger LFP in bats may have contributed to the better signal to noise ratio found in comparison of normalized strength differences.

### A better signal to noise ratio in bats leads to lower resource cost on accurate stimuli representation

The bat auditory cortex may have been more readily primed for accurate processing due to the higher signal to noise ratio, shown through CWT and model fit analyses. Forward suppression has been shown to sharpen cortical response and reduce spike rate per echo in echolocating bats (Macias et al., 2022), a feature that may also improve accuracy in call discrimination. In the comparison of signal traces at ~5 and ~40 Hz (Figure 2), bat normalized cortical activity had less jitter around stimulus response. That is, the pre-stimulus baseline was closer to its relative 0 and cortical activity adapted back closer to relative 0 after consecutive stimulus responses. Mouse normalized activity was more variable. In the ~40 Hz condition of the model fit analysis (Figure 3) bats had a higher intercept (first observed peak amplitude) in the AVREC and thalamic input layers. They subsequently had a deeper suppression of response amplitude to consecutive responses, reflected in the higher dynamic range parameter. Due to the deeper forward suppression in bats, the response amplitude generally adapted slower to stimuli in these traces at ~40 Hz, reflected in the lower rate of decay parameter in the AVREC and layer V. Mice had a weaker onset response and a shallower rate of forward suppression to consecutive responses due to higher noise in the signal trace.

Further evidence for a better signal to noise ratio in bats was provided by the normalized power scalograms from the CWT analysis (Figure 4). The bat spectral power was significantly stronger at the timepoint of stimulus onset response throughout the layers. However, the background spectral power was significantly stronger in mice, especially at higher oscillation frequencies where stimulus response was at a shorter time scale. The better signal to noise ratio found in bats may allow stimulus processing to be more temporally precise at a relatively lower metabolic cost to the neural populations.

### Phase coherence revealed lower inter-trial variability in bat auditory response profiles

The phase coherence scalograms from CWT analysis revealed a significantly stronger inter-trial broadband phase coherence at the time of stimulus and following response (Figure 5). We had expected mice to have better coherence, the probes being chronically implanted in one site rather than moving around the penetration site per measurement (as in the bat dataset). Counterintuitively, the mouse data had consistently greater variability between measurements. This was notable in the level of background jitter in the AVREC and layer traces in mice and that their higher frequency stimulus presentation peak amplitudes were less aligned with the exponential model fits compared to bats. The difference in group size may contribute partially to this discrepancy, with more measurements (n=46) in the bat dataset than in the mouse dataset (n=28), creating a greater smoothing in the averages. However, when non-normalized individual animal data is considered (Supp Figures 1-3), the signals still maintain the same features relevant to analysis within individuals. Therefore, the difference in neuronal variability appeared to be a species effect, not a group size effect.

Using two independent datasets for comparison innately bears the challenge to handle variability introduced by different stress levels between both species based on head-fixation techniques, session lengths, or the differences of the stimulus class, which may affect our physiological results. However, bat specialization in auditory perception also likely contributes to the significant discrepancies in the described cortical response variability. Bats require temporal precision in their echolocation and communication calls for behavioral success. That a bat has more accurate and less variable auditory responses to consecutive stimuli than a mouse, is evidence of successful specialization of shared architecture for different behaviors. Our analysis approach may foster further cross-species comparisons that could give us insight into the differential ways the mammalian cortex introduces or limits variability in populations of neurons based on ecological need.

### Phase amplitude coupling fundamentally different between species

PAC is a well-established phenomenon throughout the brain and neocortex (Esghaei et al., 2015; Helfrich & Knight, 2016; Lisman & Jensen, 2013; O’Connell et al., 2015; Sotero et al., 2015; Spaak et al., 2012; Xiao et al., 2019), and has been implicated in a variety of relevant functional tasks such as interareal communication and information binding (Colgin et al., 2009; Daume et al., 2017) and in information transfer across neural tissue (Bonnefond et al., 2017; Gourévitch et al., 2020). The functional use of PAC for information binding or the segmentation of continuous stimuli into slower times-scales of perceptual units, may be conserved through evolution as a shared mechanism in mammals (Garcia-Rosales et al., 2020). For example, in humans, theta gamma PAC has been implicated in efficient processing of speech phenomes into words and sentences (Gross et al., 2013; Lizarazu et al., 2019; Zion Golumbic et al., 2013). García-Rosales et al. (2020) suggested that bats could utilize this parsing strategy on echolocation to make sense of their auditory, and therefore spatial, scene. We ran a PAC coupling analysis on the A1 datasets for bats and mice at low and high stimulus presentation conditions and during spontaneous activity.

PAC in mice was similar in both stimulation frequencies, strongest in layer III/IV, but weakest during spontaneous activity. Coupling therefore seemed to depend on a stimulus being present but was not sensitive to the frequency of presentation (Figure 6 & 7). By contrast, bat PAC was different for low and high frequency stimuli and for spontaneous activity. Especially in layer III/IV, PAC was stronger at 5.28 Hz and strongest during spontaneous activity, indicating a stimulus dependent coupling strength.

Bats and mice were most different in their spontaneous activity PAC, where bats had a significantly stronger and broader area of coupling. Several studies have found that coupling may assist remote activity across neuronal assemblies in the absence of a current stimuli to process (Wang et al., 2012; Weaver et al., 2016). This may explain the stronger spontaneous vs stimulus response PAC in bats but not the weaker spontaneous vs stimulus response PAC in mice. Regardless of signal frequency, or whether it was stimulus derived or during resting-state, coupling was centered around delta/high gamma in mice and theta/low gamma in bats. The similarity in the bat PAC with human speech perception theta gamma coupling may support the hypothesized auditory scene parsing (García-Rosales et al., 2020; Gross et al., 2013; Lizarazu et al., 2019; Zion Golumbic et al., 2013). However, this region may have had greater local A1 coupling because the stimuli was specifically a bat vocalization syllable. It is also possible that the delta/high gamma coupling in mice supports the same task in a different temporal scale.

## Conclusion

When comparing two fundamentally different species, with alien subjective experiences, an analysis like this cannot say more than that these species have different interpretations of objective, external sound waves (Nagel, 1974). Nevertheless, cross-species comparisons can serve as valuable framework in consideration of shared, convergent, and divergent evolutionary adaptation (Sherry, 2007). Seba’s short-tailed bats have adapted to an ecological niche which requires accurate temporal auditory perception during 3-dimensional navigation in flight and complex social communication. For mice accurate sound representation may be less fundamental to find behavioral success in their environment. In alignment with this hypothesis, we have found that the neuronal signature of the auditory cortex in bats shows a significantly better signal to noise ratio, more accurate and less variable following responses to consecutive stimuli, far higher inter-trial phase coherence, and fundamentally different PAC profiles compared to mice. These discrepancies likely do not stem from differing cortical architecture alone. Rather, findings here implicate evolutionary divergence in the recruitment of mammalian auditory cortical microcircuitry.

Further research should test other possible causes for observed discrepancies, including if they were at least partially stimulus driven, if the size of pyramidal neurons in the cortex were larger in bats, or if bats have a higher ratio of silent neurons or synapses than mice (Ovsepian, 2019; Shoham et al., 2006). The latter may explain where cortical variability, a necessary feature for fast adaptation, is being reserved in bats and also why their background activity was lower compared to stimulus response.

## Additional Information

### Acronyms

A1 – primary auditory cortex

AVREC – average rectified (CSD)

CSD – current source density

CWT – continuous wavelet transform

I/II – supragranular layers 1 and 2

III/IV – granular layers 3 and 4

mwu – Mann-Whitney U (test)

PAC – phase amplitude coupling

V – infragranular layer 5

V – infragranular layer 6

zMI – z-scored modulation index

### Author contributions

Experiments were performed in the laboratories of the Leibniz-Institute for Neurobiology, Magdeburg (Germany), by KED and Institute for Cell Biology and Neuroscience, Goethe-University, Frankfurt/M (Germany) by FGR. Study was designed by KED, FGR, JCH, and MFKH. JCH and MFKH supervised the project. Data and statistical analysis were performed by KED and RK with input from FGR. KED prepared the figures and wrote the initial paper draft. Manuscript was edited by KED, FGR, RK, JCH, and MFKH. All authors reviewed the manuscript, approve of the final version, and agree to be accountable for all aspects of the work. All persons designated as authors qualify and all who qualify are listed as authors.

### Data Availability

Scripts and data used in this research will be available from time of peer-reviewed publication at https://github.com/CortXplorer and https://figshare.com/ or on request.

## Acknowledgements

We would like to thank Dr. Michael Lippert and Alisa Vlasenko for their assistance with the head-fixation setup and the surgery for mice and Gisa Prange for help with the histology for bats.

## Supplemental Figures

**Supp Figure 1.**
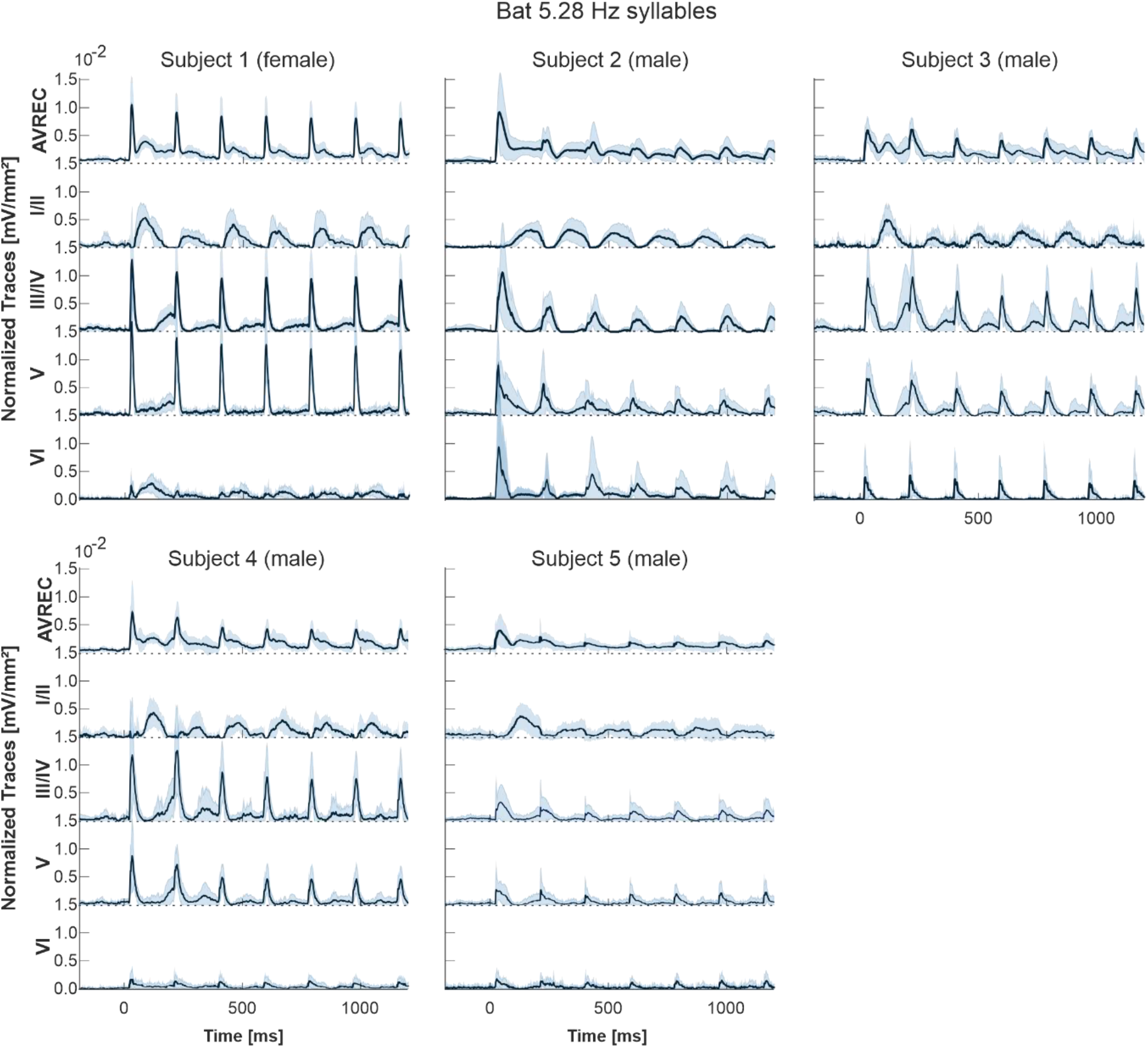
AVREC and layer traces for bat subjects at 5.28 Hz. Bat averaged auditory cortex AVREC trace (top) and all layer traces (I/II, III/IV, V, VI in descending order) per subject (n=5, 6, 8, 9, and 18 respectively), in response to 5.28 Hz click-like distress calls. Layer traces were calculated on sink activity only. Confidence intervals are shown in SEM. Traces were not normalized here but were normalized for the group average and further analysis; normalization was per measurement according to the first detected peak of the AVREC at 2 Hz (not shown).

**Supp Figure 2.**
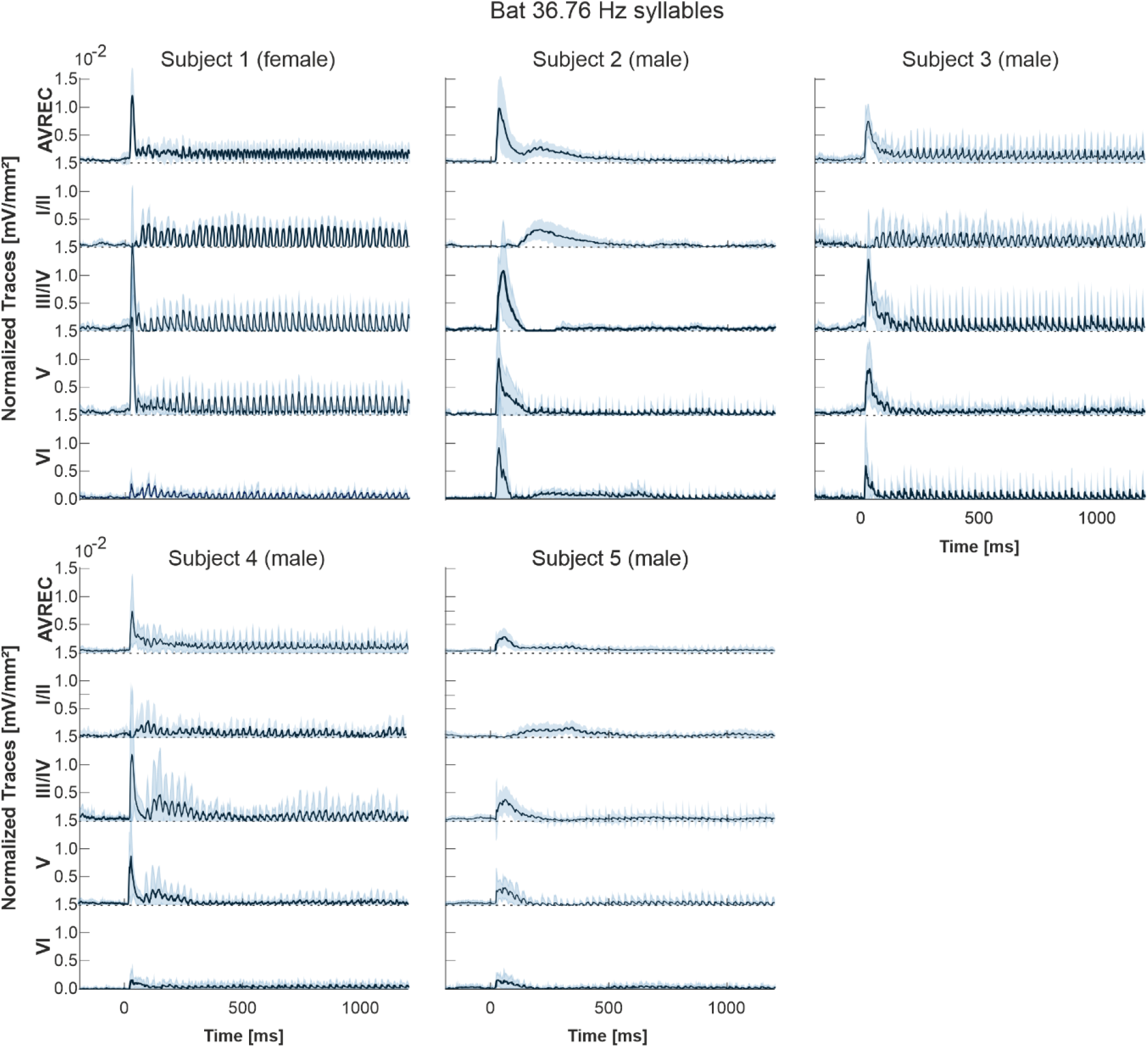
AVREC and layer traces for bat subjects at 36.76 Hz. Bat averaged auditory cortex AVREC trace (top) and all layer traces (I/II, III/IV, V, VI in descending order) per subject (n=5, 6, 8, 9, and 18 respectively), in response to 36.76 Hz click-like distress calls. Layer traces were calculated on sink activity only. Confidence intervals are shown in SEM. Traces were not normalized here but were normalized for the group average and further analysis; normalization was per measurement according to the first detected peak of the AVREC at 2 Hz (not shown).

**Supp Figure 3.**
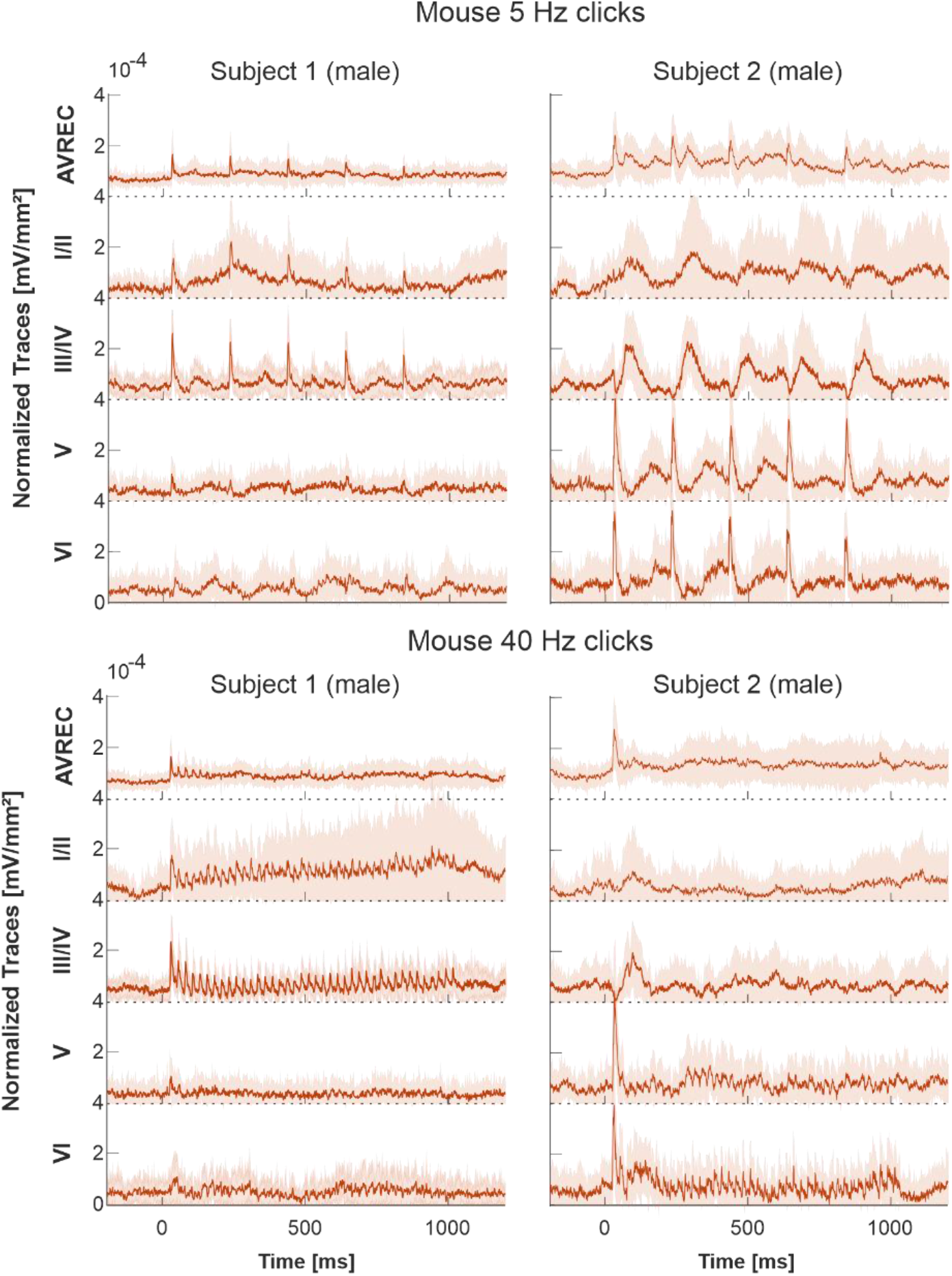
AVREC and layer traces for mouse subjects. Mouse averaged auditory cortex AVREC trace (top) and all layer traces (I/II, III/IV, V, VI in descending order) per subject (n=14 and 14 respectively), in response to 5 and 40 Hz click-trains. Layer traces were calculated on sink activity only. Confidence intervals are shown in SEM. Traces were not normalized here but were normalized for the group average and further analysis; normalization was per measurement according to the first detected peak of the AVREC at 2 Hz (not shown).

**Supp Figure 4.**
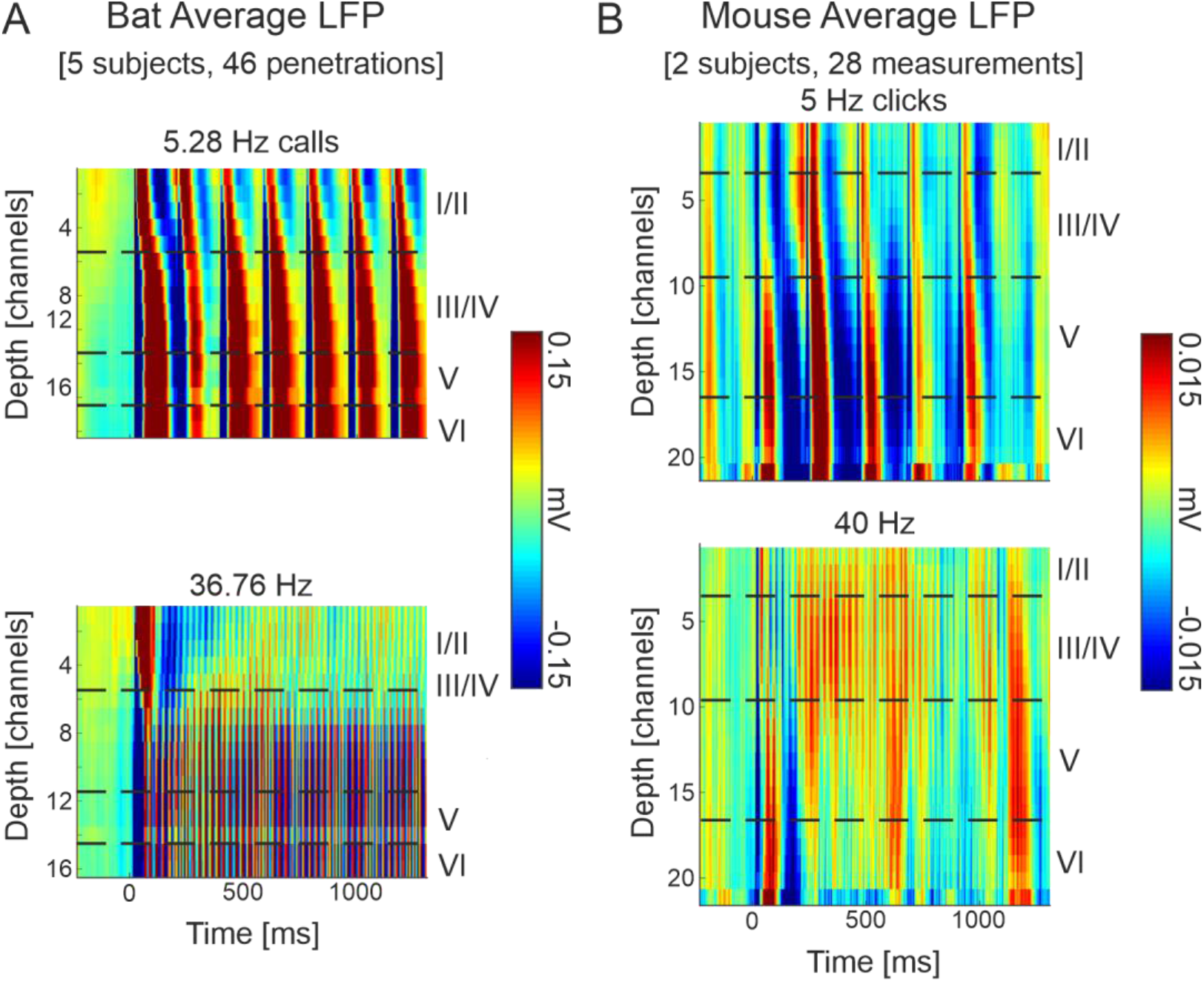
Grand average local field potential profiles. **A:** Seba’s short-tailed bats (n = 46) grand averaged cortical response to a click-like distress call presented repetitively at 5.28 Hz (top) and 36.76 Hz (bottom). **B:** C57/B6 mice (n = 28) grand averaged cortical response to a click train presented at 5 Hz (top) and 40 Hz (bottom). The LFP profiles show the pattern of temporal processing (ms) within the cortical depth (channels are 50 µm apart). Representative layer assignment is indicated with horizontal dashed lines based on CSDs (Figure 1). Current sinks (blue), represent areas of excitatory synaptic population activity. Note the different c-axis scales: with much stronger signal from bats, and the different depth scales: slightly thicker cortex for mice, ~20 channels or ~1 mm, than bats 16 channels or ~750 µm.

**Supp Figure 5.**
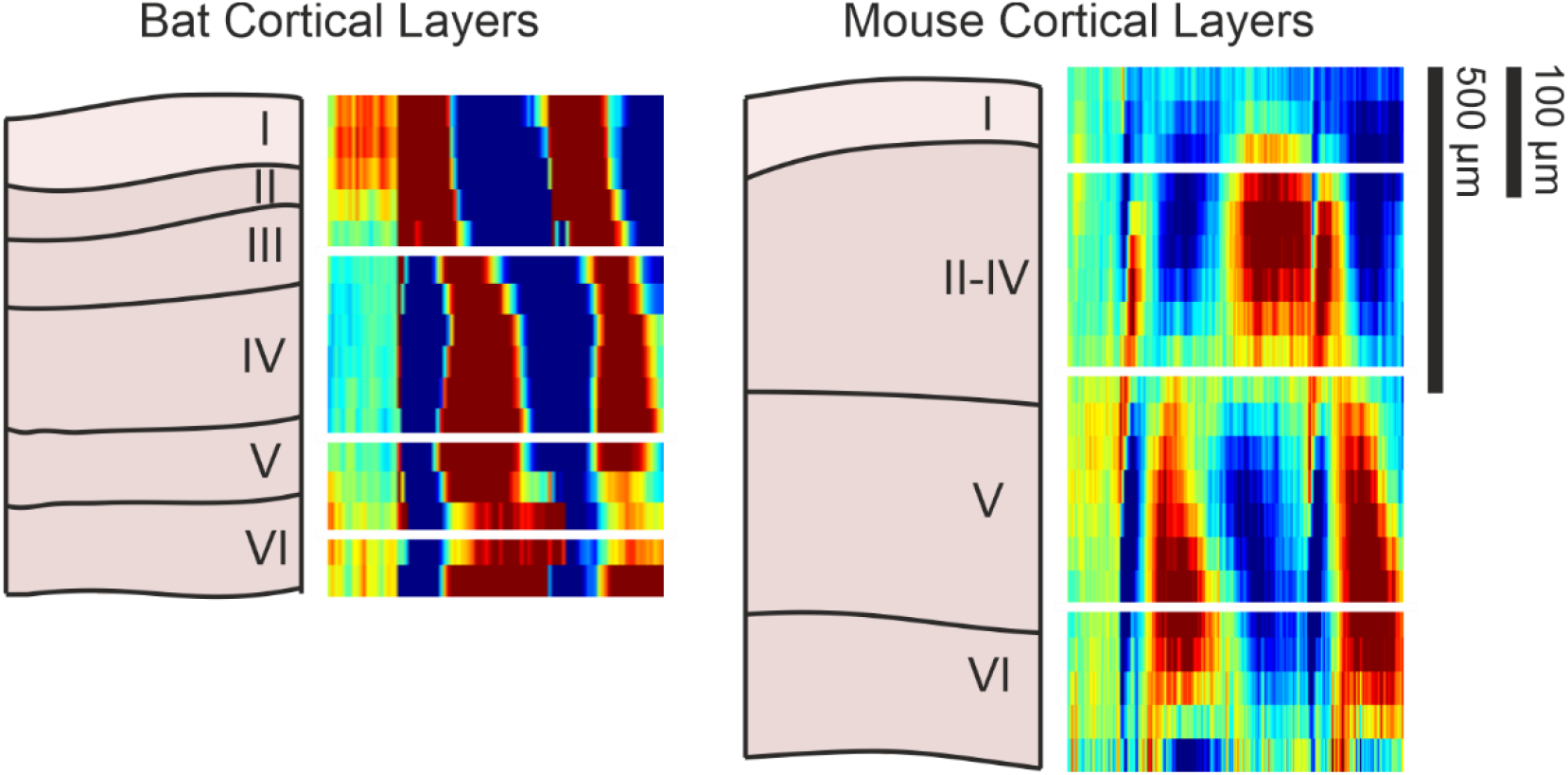
Anatomical comparison; A representation of an A1 columnal section and lines denoting rough layer boundaries were generated based on figures from García-Rosales et al. (2019) for Seba’s short-tailed bat A1 laminar anatomy and from Chang & Kawai (2018) for black 6 mouse A1 laminar anatomy. These are set next to appropriately and relatively sized group average CSD profiles (Figure 1; 1 mm for mice and 750 µm for bats) to show representative layer designations in comparison with these references.

**Supp Figure 6.**
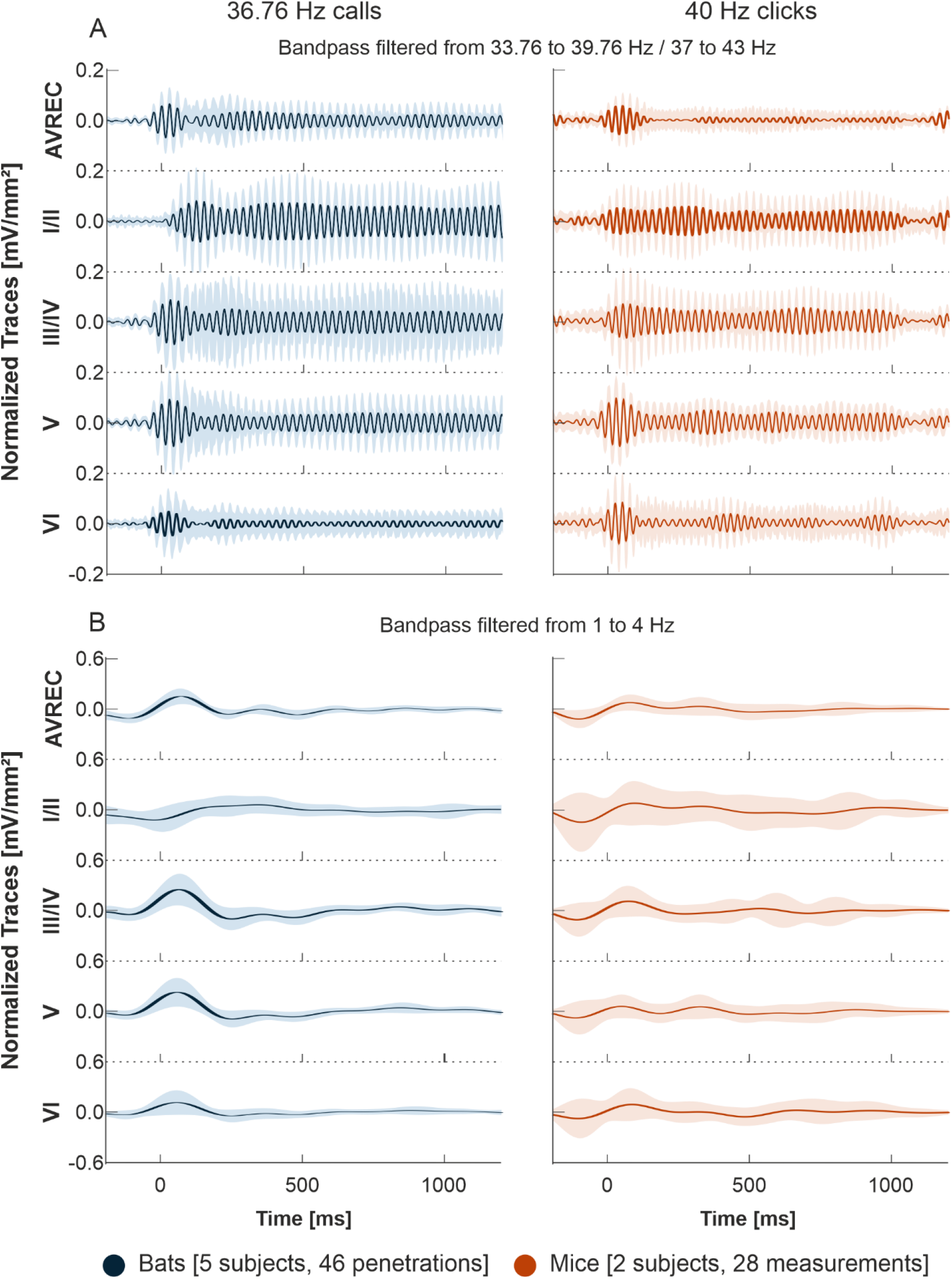
AVREC and layer traces bandpass filtered. **A:** Bat (n=46) averaged auditory cortex AVREC trace (top) and all layer traces (I/II, III/IV, V, VI in descending order), in response to 36.76 Hz click-like distress calls (blue) bandpass filtered 3 Hz above and below presentation frequency and **C:** bandpass filtered from 1 to 4 Hz. **B:** Mouse (n=28) averaged auditory cortex AVREC and layer traces, in response to 40 Hz click-trains (orange) bandpass filtered 3 Hz above and below presentation frequency and **D:** bandpass filtered from 1 to 4 Hz. Layer traces were calculated on sink activity only. Confidence intervals are shown in SEM. Traces were all normalized per measurement (bat n=46, mouse n=28). Normalization was done by dividing traces by the first detected peak of the AVREC at 2 Hz (not shown).

**Supp Figure 7.**
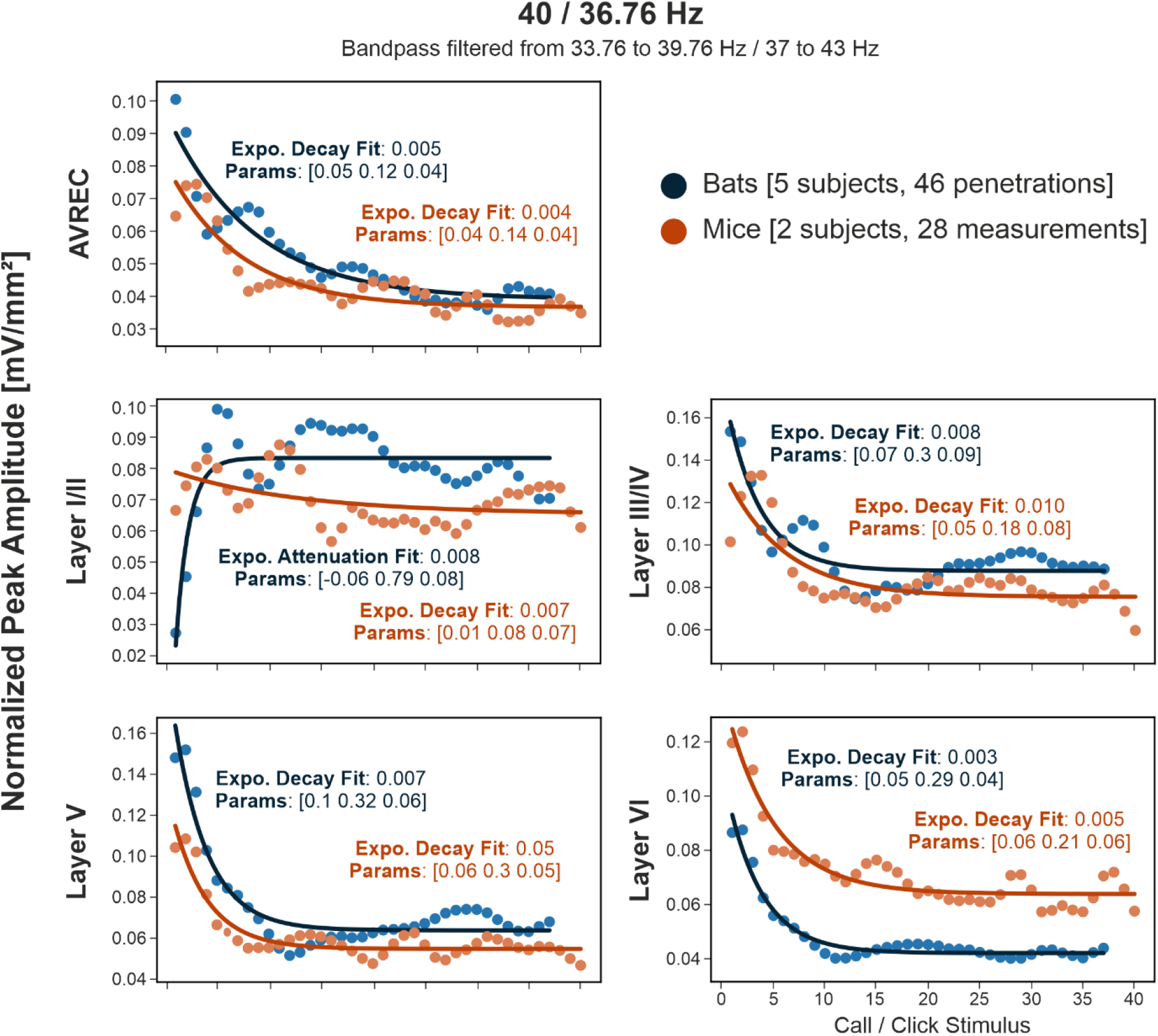
Model Fit Analysis bandpass filtered. Bandpass filtered +/- 3 Hz around the presentation frequency, bat (blue, n=46) and mouse (orange, n=28) group-averaged response peak amplitudes over consecutive stimulus repetition of 40 or 36.76 Hz with overlaid model fit. The model selected, exponential or linear decay is overlaid, along with the fit value calculated by RMSE and the model parameters. The closer to zero that the model fit is, the better. For expo.: parameters are [dynamic range, rate of decay, offset]. For linear (not present): [slope, intercept]

## References

Beetz, M. J., Kordes, S., García-Rosales, F., Kössl, M., & Hechavarría, J. C. (2017). Processing of Natural Echolocation Sequences in the Inferior Colliculus of Seba’s Fruit Eating Bat, Carollia perspicillata. ENeuro, 4(6), ENEURO.0314-17.2017. https://doi.org/10.1523/ENEURO.0314-17.2017

Bonnefond, M., Kastner, S., & Jensen, O. (2017). Communication between Brain Areas Based on Nested Oscillations. ENeuro, 4(2), ENEURO.0153-16.2017. https://doi.org/10.1523/ENEURO.0153-16.2017

Brunk, M. G. K., Deane, K. E., Kisse, M., Deliano, M., Vieweg, S., Ohl, F. W., Lippert, M. T., & Happel, M. F. K. (2019). Optogenetic stimulation of the VTA modulates a frequency-specific gain of thalamocortical inputs in infragranular layers of the auditory cortex. Scientific Reports, 9(1), 20385. https://doi.org/10.1038/s41598-019-56926-6

Buzsáki, G., Anastassiou, C. A., & Koch, C. (2012). The origin of extracellular fields and currents—EEG, ECoG, LFP and spikes. Nature Reviews Neuroscience, 13(6), 407–420. https://doi.org/10.1038/nrn3241

Cardillo, G. (2009). *MWWTEST: Mann-Whitney-Wilcoxon non parametric test for two unpaired samples.* http://www.mathworks.com/matlabcentral/fileexchange/25830

Chang, M., & Kawai, H. D. (2018). A characterization of laminar architecture in mouse primary auditory cortex. Brain Structure and Function, 223(9), 4187–4209. https://doi.org/10.1007/s00429-018-1744-8

Chen, G., Zhang, Y., Li, X., Zhao, X., Ye, Q., Lin, Y., Tao, H. W., Rasch, M. J., & Zhang, X. (2017). Distinct Inhibitory Circuits Orchestrate Cortical beta and gamma Band Oscillations. Neuron, 96(6), 1403–1418.e6. https://doi.org/10.1016/j.neuron.2017.11.033

Colgin, L. L., Denninger, T., Fyhn, M., Hafting, T., Bonnevie, T., Jensen, O., Moser, M.-B., & Moser, E. I. (2009). Frequency of gamma oscillations routes flow of information in the hippocampus. Nature, 462(7271), Article 7271. https://doi.org/10.1038/nature08573

Daume, J., Gruber, T., Engel, A. K., & Friese, U. (2017). Phase-Amplitude Coupling and Long-Range Phase Synchronization Reveal Frontotemporal Interactions during Visual Working Memory. Journal of Neuroscience, 37(2), 313–322. https://doi.org/10.1523/JNEUROSCI.2130-16.2016

De Franceschi, G., & Barkat, T. R. (2021). Task-induced modulations of neuronal activity along the auditory pathway. Cell Reports, 37(11), 110115. https://doi.org/10.1016/j.celrep.2021.110115

Deane, K. E., Brunk, M. G. K., Curran, A. W., Zempeltzi, M. M., Ma, J., Lin, X., Abela, F., Aksit, S., Deliano, M., Ohl, F. W., & Happel, M. F. K. (2020). Ketamine anaesthesia induces gain enhancement via recurrent excitation in granular input layers of the auditory cortex. Journal of Physiology, 598(13), 2741–2755. https://doi.org/10.1113/JP279705

Deane, K. E., Klymentiev, R., Heck, J., Mark, M. D., Ohl, F. W., Heine, M., & Happel, M. F. K. (2022). Inhibiting presynaptic calcium channel mobility in the auditory cortex suppresses synchronized input processing (p. 2022.03.30.486338). bioRxiv. https://doi.org/10.1101/2022.03.30.486338

Esghaei, M., Daliri, M. R., & Treue, S. (2015). Attention Decreases Phase-Amplitude Coupling, Enhancing Stimulus Discriminability in Cortical Area MT. Frontiers in Neural Circuits, 9. https://www.frontiersin.org/article/10.3389/fncir.2015.00082

Fonseca, A. H., Santana, G. M., Bosque Ortiz, G. M., Bampi, S., & Dietrich, M. O. (2021). Analysis of ultrasonic vocalizations from mice using computer vision and machine learning. ELife, 10, e59161. https://doi.org/10.7554/eLife.59161

García-Rosales, F., López-Jury, L., González-Palomares, E., Cabral-Calderín, Y., Kössl, M., & Hechavarria, J. C. (2020). Phase-amplitude coupling profiles differ in frontal and auditory cortices of bats. European Journal of Neuroscience, n/a(n/a). https://doi.org/10.1111/ejn.14986

García-Rosales, F., López-Jury, L., González-Palomares, E., Wetekam, J., Cabral-Calderín, Y., Kiai, A., Kössl, M., & Hechavarría, J. C. (2022). Echolocation-related reversal of information flow in a cortical vocalization network. Nature Communications, 13(1), Article 1. https://doi.org/10.1038/s41467-022-31230-6

García-Rosales, F., Röhrig, D., Weineck, K., Röhm, M., Lin, Y.-H., Cabral-Calderin, Y., Kössl, M., & Hechavarria, J. C. (2019). Laminar specificity of oscillatory coherence in the auditory cortex. Brain Structure and Function, 224(8), 2907–2924. https://doi.org/10.1007/s00429-019-01944-3

Gire, D. H., Kapoor, V., Arrighi-Allisan, A., Seminara, A., & Murthy, V. N. (2016). Mice develop efficient strategies for foraging and navigation using complex natural stimuli. Current Biology : CB, 26(10), 1261–1273. https://doi.org/10.1016/j.cub.2016.03.040

Givre, S. J. J., Schroeder, C. E. E., & Arezzo, J. C. C. (1994). Contribution of extrastriate area V4 to the surface-recorded flash VEP in the awake macaque. Vision Research, 34(4), 415–428. https://doi.org/10.1016/0042-6989(94)90156-2

Gourévitch, B., Martin, C., Postal, O., & Eggermont, J. J. (2020). Oscillations in the auditory system and their possible role. Neuroscience and Biobehavioral Reviews, 113(April), 507–528. https://doi.org/10.1016/j.neubiorev.2020.03.030

Groppe, D. M., Urbach, T. P., & Kutas, M. (2011). Mass univariate analysis of event-related brain potentials/fields I: A critical tutorial review. In Psychophysiology (Vol. 48, Issue 12, pp. 1711–1725). Blackwell Publishing Inc. https://doi.org/10.1111/j.1469-8986.2011.01273.x

Gross, J., Hoogenboom, N., Thut, G., Schyns, P., Panzeri, S., Belin, P., & Garrod, S. (2013). Speech Rhythms and Multiplexed Oscillatory Sensory Coding in the Human Brain. PLOS Biology, 11(12), e1001752. https://doi.org/10.1371/journal.pbio.1001752

Grundy, D. (2015). Principles and standards for reporting animal experiments in The Journal of Physiology and Experimental Physiology. In Journal of Physiology (Vol. 593, Issue 12, pp. 2547–2549). Blackwell Publishing Ltd. https://doi.org/10.1113/JP270818

Hagemann, C., Vater, M., & Kössl, M. (2011). Comparison of properties of cortical echo delay-tuning in the short-tailed fruit bat and the mustached bat. Journal of Comparative Physiology A, 197(5), 605–613. https://doi.org/10.1007/s00359-010-0530-8

Happel, M. F. K., Deliano, M., Handschuh, J., & Ohl, F. W. (2014). Dopamine-Modulated Recurrent Corticoefferent Feedback in Primary Sensory Cortex Promotes Detection of Behaviorally Relevant Stimuli. Journal of Neuroscience, 34(4), 1234–1247. https://doi.org/10.1523/JNEUROSCI.1990-13.2014

Happel, M. F. K., Jeschke, M., & Ohl, F. W. (2010). Spectral Integration in Primary Auditory Cortex Attributable to Temporally Precise Convergence of Thalamocortical and Intracortical Input. The Journal of Neuroscience, 30(33), 11114–11127.

Happel, M. F. K., & Ohl, F. W. (2017). Compensating Level-Dependent Frequency Representation in Auditory Cortex by Synaptic Integration of Corticocortical Input. PLOS ONE, 12(1), e0169461. https://doi.org/10.1371/journal.pone.0169461

Hashikawa, T., Molinari, M., Rausell, E., & Jones, E. G. (1995). Patchy and laminar terminations of medial geniculate axons in monkey auditory cortex. The Journal of Comparative Neurology, 362(2), 195–208. https://doi.org/10.1002/cne.903620204

Hechavarría, J. C., Macías, S., Vater, M., Mora, E. C., & Kössl, M. (2013). Evolution of neuronal mechanisms for echolocation: Specializations for target-range computation in bats of the genus Pteronotus. The Journal of the Acoustical Society of America, 133(1), 570–578. https://doi.org/10.1121/1.4768794

Hechavarría, J. C., Macías, S., Vater, M., Voss, C., Mora, E. C., & Kössl, M. (2013). Blurry topography for precise target-distance computations in the auditory cortex of echolocating bats. Nature Communications, 4, 2587. https://doi.org/10.1038/ncomms3587

Helfrich, R. F., & Knight, R. T. (2016). Oscillatory Dynamics of Prefrontal Cognitive Control. Trends in Cognitive Sciences, 20(12), 916–930. https://doi.org/10.1016/j.tics.2016.09.007

Herculano-Houzel, S., Collins, C. E., Wong, P., & Kaas, J. H. (2007). Cellular scaling rules for primate brains. Proceedings of the National Academy of Sciences, 104(9), 3562–3567. https://doi.org/10.1073/pnas.0611396104

Hoglen, N. E. G., Larimer, P., Phillips, E. A. K., Malone, B. J., & Hasenstaub, A. R. (2018). Amplitude modulation coding in awake mice and squirrel monkeys. Journal of Neurophysiology, 119(5), 1753–1766. https://doi.org/10.1152/jn.00101.2017

Kanwal, J. S. (2012). Right–left asymmetry in the cortical processing of sounds for social communication vs. Navigation in mustached bats. European Journal of Neuroscience, 35(2), 257–270. https://doi.org/10.1111/j.1460-9568.2011.07951.x

Kanwal, J. S., & Rauschecker, J. P. (2007). Auditory cortex of bats and primates: Managing species-specific calls for social communication. Frontiers in Bioscience : A Journal and Virtual Library, 12, 4621–4640.

Karmos, G., Lakatos, P., Pincze, Z., Rajkai, C., & Ulbert, I. (2002). Frequency of gamma activity is modulated by motivation in the auditory cortex of cat. Acta Biologica Hungarica, 53(4), 473–483. https://doi.org/10.1556/ABiol.53.2002.4.8

Kato, H. K., Gillet, S. N., & Isaacson, J. S. (2015). Flexible Sensory Representations in Auditory Cortex Driven by Behavioral Relevance. Neuron, 88(5), 1027–1039. https://doi.org/10.1016/j.neuron.2015.10.024

Kikuchi, Y., Attaheri, A., Wilson, B., Rhone, A. E., Nourski, K. V., Gander, P. E., Kovach, C. K., Kawasaki, H., Griffiths, T. D., Howard, M. A., & Petkov, C. I. (2017). Sequence learning modulates neural responses and oscillatory coupling in human and monkey auditory cortex. PLoS Biology, 15(4), e2000219. https://doi.org/10.1371/journal.pbio.2000219

Lachaux, J.-P., Rodriguez, E., Martinerie, J., & Varela, F. J. (1999). Measuring phase synchrony in brain signals. Human Brain Mapping, 8(4), 194–208. https://doi.org/10.1002/(SICI)1097-0193(1999)8:4<194::AID-HBM4>3.0.CO;2-C

Lakatos, P., Szilágyi, N., Pincze, Z., Rajkai, C., Ulbert, I., & Karmos, G. (2004). Attention and arousal related modulation of spontaneous gamma-activity in the auditory cortex of the cat. Brain Research. Cognitive Brain Research, 19(1), 1–9. https://doi.org/10.1016/j.cogbrainres.2003.10.023

Lilly, J. M., & Olhede, S. C. (2012). Generalized morse wavelets as a superfamily of analytic wavelets. IEEE Transactions on Signal Processing, 60(11), 6036–6041. https://doi.org/10.1109/TSP.2012.2210890

Linden, J. F., & Schreiner, C. E. (2003). Columnar Transformations in Auditory Cortex? A Comparison to Visual and Somatosensory Cortices. Cerebral Cortex, 13(1), 83–89. https://doi.org/10.1093/cercor/13.1.83

Lisman, J. E., & Jensen, O. (2013). The Theta-Gamma Neural Code. In Neuron (Vol. 77). https://doi.org/10.1016/j.neuron.2013.03.007

Lizarazu, M., Lallier, M., & Molinaro, N. (2019). Phase−amplitude coupling between theta and gamma oscillations adapts to speech rate. Annals of the New York Academy of Sciences, 1453(1), 140–152. https://doi.org/10.1111/nyas.14099

López-Jury, L., García-Rosales, F., González-Palomares, E., Kössl, M., & Hechavarria, J. C. (2021). Acoustic Context Modulates Natural Sound Discrimination in Auditory Cortex through Frequency-Specific Adaptation. Journal of Neuroscience, 41(50), 10261–10277. https://doi.org/10.1523/JNEUROSCI.0873-21.2021

Lu, Z.-L., Williamson, S., & Kaufman, L. (1993). Lu ZL, Williamson SJ, Kaufman L. Behavioral lifetime of human auditory sensory memory predicted by physiological measures. Science 258: 1668-1670. Science (New York, N.Y.), 258, 1668–1670. https://doi.org/10.1126/science.1455246

MacDonald, K. D., & Barth, D. S. (1995). High frequency (gamma-band) oscillating potentials in rat somatosensory and auditory cortex. Brain Research, 694(1–2), 1–12. https://doi.org/10.1016/0006-8993(95)00662-a

Macias, S., Bakshi, K., & Smotherman, M. (2022). Faster Repetition Rate Sharpens the Cortical Representation of Echo Streams in Echolocating Bats. ENeuro, 9(1). https://doi.org/10.1523/ENEURO.0410-21.2021

Maris, E., Schoffelen, J.-M., & Fries, P. (2007). Nonparametric statistical testing of coherence differences. Journal of Neuroscience Methods, 163(1), 161–175. https://doi.org/10.1016/j.jneumeth.2007.02.011

McMullen, N. T., & Glaser, E. M. (1982). Morphology and laminar distribution of nonpyramidal neurons in the auditory cortex of the rabbit. The Journal of Comparative Neurology, 208(1), 85–106. https://doi.org/10.1002/cne.902080107

Mitzdorf, U. (1985). Current source-density method and application in cat cerebral cortex: Investigation of evoked potentials and EEG phenomena. PHYSIOLOGICAL REVIEWS, 65, 64.

Mountcastle, V. B. (1997). The columnar organization of the neocortex. Brain, 120(4), 701–722. https://doi.org/10.1093/brain/120.4.701

Muramatsu, S., Toda, M., Nishikawa, J., & Tateno, T. (2019). Sound- and current-driven laminar profiles and their application method mimicking acoustic responses in the mouse auditory cortex in vivo. Brain Research, 1721, 146312. https://doi.org/10.1016/j.brainres.2019.146312

Nagel, T. (1974). What Is It Like to Be a Bat? The Philosophical Review, 83(4), 435. https://doi.org/10.2307/2183914

Neophytou, D., Arribas, D. M., Arora, T., Levy, R. B., Park, I. M., & Oviedo, H. V. (2022). Differences in temporal processing speeds between the right and left auditory cortex reflect the strength of recurrent synaptic connectivity. PLOS Biology, 20(10), e3001803. https://doi.org/10.1371/journal.pbio.3001803

O’Connell, M. N., Barczak, A., Ross, D., McGinnis, T., Schroeder, C. E., & Lakatos, P. (2015). Multi-Scale Entrainment of Coupled Neuronal Oscillations in Primary Auditory Cortex. Frontiers in Human Neuroscience, 9. https://www.frontiersin.org/article/10.3389/fnhum.2015.00655

Olhede, S. C., & Walden, A. T. (2002). Generalized Morse wavelets. IEEE Transactions on Signal Processing, 50(11), 2661–2670. https://doi.org/10.1109/TSP.2002.804066

Ovsepian, S. V. (2019). The dark matter of the brain. Brain Structure and Function, 224(3). https://doi.org/10.1007/s00429-019-01835-7

Schaefer, M. K., Hechavarría, J. C., & Kössl, M. (2015). Quantification of mid and late evoked sinks in laminar current source density profiles of columns in the primary auditory cortex. Frontiers in Neural Circuits, 9, 1–16. https://doi.org/10.3389/fncir.2015.00052

Schroeder, C. E., Mehta, A. D., & Givre, S. J. (1998). A spatiotemporal profile of visual system activation revealed by current source density analysis in the awake macaque. Cerebral Cortex, 8(7), 575–592. https://doi.org/10.1093/cercor/8.7.575

Shahriari, Y., Krusienski, D., Dadi, Y. S., Seo, M., Shin, H.-S., & Choi, J. H. (2016). Impaired auditory evoked potentials and oscillations in frontal and auditory cortex of a schizophrenia mouse model. The World Journal of Biological Psychiatry: The Official Journal of the World Federation of Societies of Biological Psychiatry, 17(6), 439–448. https://doi.org/10.3109/15622975.2015.1112036

Sherry, D. F. (2007). Cross-Species Comparisons. In Ciba Foundation Symposium 208—Characterizing Human Psychological Adaptations (pp. 181–194). John Wiley & Sons, Ltd. https://doi.org/10.1002/9780470515372.ch10

Shoham, S., O’Connor, D. H., & Segev, R. (2006). How silent is the brain: Is there a “dark matter” problem in neuroscience? Journal of Comparative Physiology A, 192(8), 777–784. https://doi.org/10.1007/s00359-006-0117-6

Sotero, R. C., Bortel, A., Naaman, S., Mocanu, V. M., Kropf, P., Villeneuve, M. Y., & Shmuel, A. (2015). Laminar Distribution of Phase-Amplitude Coupling of Spontaneous Current Sources and Sinks. Frontiers in Neuroscience, 9. https://www.frontiersin.org/article/10.3389/fnins.2015.00454

Spaak, E., Bonnefond, M., Maier, A., Leopold, D. A., & Jensen, O. (2012). Layer-Specific Entrainment of Gamma-Band Neural Activity by the Alpha Rhythm in Monkey Visual Cortex. Current Biology, 22(24), 2313–2318. https://doi.org/10.1016/j.cub.2012.10.020

Szymanski, F. D., Garcia-Lazaro, J. A., & Schnupp, J. W. H. (2009). Current Source Density Profiles of Stimulus-Specific Adaptation in Rat Auditory Cortex. Journal of Neurophysiology, 102(3), 1483–1490. https://doi.org/10.1152/jn.00240.2009

Thies, W., Kalko, E. K. V., & Schnitzler, H.-U. (1998). The roles of echolocation and olfaction in two Neotropical fruit-eating bats, Carollia perspicillata and C. castanea, feeding on Piper. Behavioral Ecology and Sociobiology, 42(6), 397–409. https://doi.org/10.1007/s002650050454

Vianney-Rodrigues, P., Iancu, O. D., & Welsh, J. P. (2011). Gamma oscillations in the auditory cortex of awake rats. The European Journal of Neuroscience, 33(1), 119–129. https://doi.org/10.1111/j.1460-9568.2010.07487.x

Virtanen, P., Gommers, R., Oliphant, T. E., Haberland, M., Reddy, T., Cournapeau, D., Burovski, E., Peterson, P., Weckesser, W., Bright, J., van der Walt, S. J., Brett, M., Wilson, J., Millman, K. J., Mayorov, N., Nelson, A. R. J., Jones, E., Kern, R., Larson, E., … Vázquez-Baeza, Y. (2020). SciPy 1.0: Fundamental algorithms for scientific computing in Python. Nature Methods, 17(3), 261–272. https://doi.org/10.1038/s41592-019-0686-2

Weineck, K., García-Rosales, F., & Hechavarría, J. C. (2020). Neural oscillations in the fronto-striatal network predict vocal output in bats. PLOS BIOLOGY, 18(3), 29. https://doi.org/10.1371/journal.pbio.3000658

Wetzel, W., Ohl, F. W., & Scheich, H. (2008). Global versus local processing of frequency-modulated tones in gerbils: An animal model of lateralized auditory cortex functions. Proceedings of the National Academy of Sciences, 105(18), 6753–6758. https://doi.org/10.1073/pnas.0707844105

Winguth, S. D., & Winer, J. A. (1986). Corticocortical connections of cat primary auditory cortex (AI): Laminar organization and identification of supragranular neurons projecting to area AII. The Journal of Comparative Neurology, 248(1), 36–56. https://doi.org/10.1002/cne.902480104

Xiao, Z., Martinez, E., Kulkarni, P. M., Zhang, Q., Hou, Q., Rosenberg, D., Talay, R., Shalot, L., Zhou, H., Wang, J., & Chen, Z. S. (2019). Cortical Pain Processing in the Rat Anterior Cingulate Cortex and Primary Somatosensory Cortex. Frontiers in Cellular Neuroscience, 13. https://www.frontiersin.org/article/10.3389/fncel.2019.00165

Yamamura, D., Sano, A., & Tateno, T. (2017). An analysis of current source density profiles activated by local stimulation in the mouse auditory cortex in vitro. Brain Research, 1659, 96–112. https://doi.org/10.1016/j.brainres.2017.01.021

Zatorre, R. J., & Belin, P. (2001). Spectral and temporal processing in human auditory cortex. Cerebral Cortex (New York, N.Y.: 1991), 11(10), 946–953. https://doi.org/10.1093/cercor/11.10.946

Zempeltzi, M. M., Kisse, M., Brunk, M. G. K., Glemser, C., Aksit, S., Deane, K. E., Maurya, S., Schneider, L., Ohl, F. W., Deliano, M., & Happel, M. F. K. (2020). Task rule and choice are reflected by layer-specific processing in rodent auditory cortical microcircuits. Communications Biology, 3(1), 345. https://doi.org/10.1038/s42003-020-1073-3

Zion Golumbic, E. M., Ding, N., Bickel, S., Lakatos, P., Schevon, C. A., McKhann, G. M., Goodman, R. R., Emerson, R., Mehta, A. D., Simon, J. Z., Poeppel, D., & Schroeder, C. E. (2013). Mechanisms Underlying Selective Neuronal Tracking of Attended Speech at a “Cocktail Party.” Neuron, 77(5), 980–991. https://doi.org/10.1016/j.neuron.2012.12.037

